# Multiple Sex Chromosome Drivers in a Mammal with Three Sex Chromosomes

**DOI:** 10.1101/2021.10.19.464942

**Authors:** Paul A. Saunders, Julie Perez, Ophélie Ronce, Frédéric Veyrunes

## Abstract

A few mammals have unusual sex determining systems whereby fertile XY females live alongside XX females and XY males. These systems are regarded as evolutionary paradoxes because of the production of sex-reversed individuals and non-viable embryos, but they nevertheless seem stable over evolutionary time. Several hypotheses have been proposed to account for their stability, including models involving sex chromosome drive (*i*.*e*., biased transmission of sex chromosomes to the next generation). Here we corroborate this hypothesis in *Mus minutoides*, a close relative of the house mouse in which the presence of XY females is due to the evolution of a third sex chromosome: a feminizing X. Through extensive molecular sexing of pups at weaning, we reveal the existence of a remarkable male sex chromosome drive system in this species, whereby direction and strength of drive is conditional upon the genotype of males’ partners: males transmit their Y to almost 80% of their offspring when mating with XX females, and only 36% when mating with XY females. Using mathematical modelling, we explore the joint evolution of these unusual sex-determining and drive systems, revealing that different sequences of events could have led to the evolution of this bizarre system, and that the “conditional” nature of sex chromosome drive stabilizes the feminizing X, and even precludes a return to a standard XX/XY system.

## Introduction

In therian mammals, sex is determined at fertilization by the X and Y chromosomes. This sex determining system evolved around 150my ago, making it one the oldest and most conserved sex determining systems known to date (1, 2). Nevertheless, a dozen mammalian species have been described with so-called unusual sex determining systems (3, 4). Among those, there are species in which fertile XY females live alongside the standard XX females and XY male. Naturally occurring XY sex-reversal has evolved at least five times independently: twice in lemmings, in the wood and collared lemmings *Myopus schisticolor* and *Dicrostonyx torquatus* (5, 6), in several species of South American field mice of the genus *Akodon* (7, 8), in the African pygmy mouse *Mus minutoides* (9), and in the Mandarin vole *Lasiopodomys mandarinus* (10, 11). In all these rodents, sex-reversal is due to a feminizing mutation on the X, rather than a loss of function of the Y. The femininizing X, generally called X*, leads to the co-existence of three female karyotypes: XX, XX* and X*Y, while all males are XY.

The evolution and maintenance of these sex determining systems (that we will refer to as polygenic systems, following (12)) has puzzled scientists for decades. In human, laboratory mice and domestic animals, male-to-female sex-reversal usually leads to a strong decrease in fertility (13), due to (i) the loss of YY embryos and (ii) the presence of a single X chromosome and the ectopic expression of Y-linked genes during meiosis, leading to increased oocyte loss (14–17). Nevertheless, in the species mentioned above, X*Y females tend to be found in high proportion (9, 18, 19). As it turns out, the monitoring of specimens in laboratory colonies revealed that the reproductive success of X*Y females is not significantly lower than that of XX and XX* females in the collared lemming (3, 20) and the South American field mouse *Akodon azarae* (21, 22), and that in the wood lemming and African pygmy mouse, X*Y females actually display enhanced breeding performances (23–25). In all cases, this absence of reduced fertility is at least in part due an increased ovulation rate (20, 21, 24, 26). Though it is easy to understand how this helps maintaining the feminizing chromosomes nowadays, it is likely that these features evolved secondarily, and that X*Y females had initially had a reduced fertility, in which case other mechanisms must have been responsible for the initial spread and maintenance of X* chromosomes.

These species share another remarkable feature, that is extremely rare among animals: sex chromosome drive. Sex chromosome drive, also called transmission distortion of sex chromosomes, is caused by selfish genetic elements that manipulate the production/function of gametes, or embryo survival, to increase their own transmission to the next generation (27–29). They have only been described in a handful of species, and are rare because of their effect on sex ratio (30). So far, sex chromosome drive has been described in four out of the five lineages with X* feminizing chromosomes. In *M. schisticolor* and *A. azarae*, X*Y females transmit their X* chromosome preferentially (X*-drive) (5, 31). It was demonstrated mathematically that this helps maintaining the X*, by increasing the frequency of the X* in the offspring of X*Y females, and reducing the proportion of YY embryos produced (32). Another type of drive *i*.*e*., male Y-drive, was identified in *D. torquatus* (33) and is suspected in *A. azarae* (31). Such drivers are expected to evolve in species with X*Y females, because they allow males to sire more sons on average, which represents an advantage as male is the rarer sex in the presence of an X* (34). Nevertheless, mathematical models shows is that Y-drive actually leads to an even more female-biased sex ratio (34, 35), because in crosses with X*Y females, Y-drive causes the production of less sons and more X*Y daughters. It was recently proposed that a solution to this problem evolved in the mandarin vole *L. mandarinus*, whereby the transmission pattern of male sex chromosomes is consistent with Y-drive in crosses with XX and XX* females, and X-drive in crosses with X*Y females, allowing the three types of females to produce more sons (11). Overall, the most commonly supported hypothesis is that sex chromosome drive evolved following the establishment of the X* chromosomes, due to selection for a balanced sex-ratio. Nevertheless, it was also proposed that the spread of feminizing mutations could be a consequence, rather than a cause, of the presence of sex chromosome drive. A first model demonstrated that Y-drive in standard XX/XY systems could favour the invasion of femininizing chromosomes because they allow to reduce the sex ratio bias induced by the former (36). More recently, it was also shown that mutant sex determiners that emerge in tight linkage with a meiotic driver will automatically increase in frequency (a form of genetic hitchhiking) (37, 38). The first model fits well with the male Y-drive observed in *D. torquatus, A. azarae* and *L. mandarinus*, and the second with the X*Y female X*-drive observed in *M. shisticolor* and *A. azarae*. Thus, there seems to be a clear link between the evolution of X* chromosomes and sex chromosome drive in rodents, but their causal connection remains ambiguous.

In an attempt to clarify the situtation, we analyzed the transmission ratio of sex chromosomes in the African pygmy mouse *M. minutoides*. By measuring sex ratio in progenies from close to 400 litters born in our pygmy mouse laboratory colony, we provide evidence for the existence of sex chromosome drive in this species: all three types of females produce litters with significantly more males than expected. Through extensive offspring genotyping, we show that the sex ratio bias is due to a strong drive of male sex chromosomes. The strength and direction of drive is dependent on female genotype: males transmit their Y much more often in crosses with XX and XX* females and their X more often in crosses with X*Y females. Building on existing models, we develop a set of analytical models to shed light on the joint evolution of male sex chromosome drive and the feminizing X* chromosome in *M. minutoides*. The originality of our approach lies in our attention to the consequences and evolution of conditional drive, whereby the bias in transmission of male X and Y depends on female genotype. We analyze how the transmission of male sex chromosomes affects the stability of the X* chromosome, show that different sequences of events could have led to the evolution of this atypical system, and finally demonstrate that the conditional nature of drive has a strong impact on the long-term persistence of the system

## Results

### Sex ratio and sex chromosome transmission

The expected sex ratio in the progenies of the XX, XX* and X*Y females, and observed sex ratio at weaning are shown in table 1. The proportion of males produced was significantly higher than expected in the three types of crosses. A test of unimodality (39) failed to detect multimodality in the distribution of mean sex-ratio for each type of female (XX females D=0.088, p-value=0.087, XX* females D=0.075, p-value=0.071, X*Y females: D=0.053, p-value=0.23), suggesting that all females produce litters with a biased sex ratio, *i*.*e*., that the genetic element(s) skewing sex ratio is (are) fixed in our captive population.

**Table 1.** Expected *vs*. observed sex ratio in the progenies of the three types of females.

| Female genotype | XX | XX* | X*Y |
| --- | --- | --- | --- |
| Expected sex ratio | 0.5 | 0.25<sup>1</sup> | 0.33<sup>1</sup> |
| Observed sex ratio<br>(overall number of offspring) | 0.79 +/- 0.13<br>(206) | 0.37 +/- 0.17<br>(370) | 0.42 +/- 0.14<br>(670) |
| Departure from expected sex ratio<br>(Binomial test) | p=<2.2e-16 | p=1.967e-07 | p=6.701e-05 |

<sup>1</sup>The expected sex ratio in the progenies of XX\* and X\*Y females is different from 0.5 because they produce viable offspring of respectively four genotypes (XX, XX\*, X\*Y and XY) and three genotypes (XX\*, X\*Y, XY).

It is straightforward that the bias in the sex ratio of the progeny of XX females results from a biased transmission of male sex chromosomes: there is an average of 79% of males in their progeny, meaning that males transmit their Y chromosome to roughly 80% of their offspring (Y-drive). It is less straightforward to determine whether sex ratio biases in the progenies of XX* and X*Y females are due to a skewed transmission of male or female sex chromosomes (or both). We therefore genotyped all of their offspring (table 2A). The transmission ratio of sex chromosomes in XX* and X*Y females was not significantly different from 50:50 (table 2B), in contrast to that of males: those paired with XX* females transmit their Y to almost 80% of their descendants (like males paired to XX females), and surprisingly, those paired to X*Y females transmit their X chromosome more often (X-drive), their Y chromosome being transmitted to only 36% of their offspring.

**Table 2.** Transmission ratios of sex chromosomes. A: results of genotyping, number of each type of offspring in the progeny of the three types of crosses. B. Transmission ratios of sex chromosomes.

| Female genotype |  | XX | XX* |  | X*Y |  |
| --- | --- | --- | --- | --- | --- | --- |
| Sex chromosomes |  | X | X | X* | X* | Y |
| Males | X | 43 | 37 | 52 | 248 | 283 |
|  | Y | 163 | 135 | 146 | 139 | † |

B.
|  | transmission ratio | p-value (binomial test) |
| --- | --- | --- |
| Males with XX females | **Y: 0.791**(95% CI: 0.730-0.845) | **<2.2e-16** |
| Males with XX* females | **Y: 0.760** (95% CI: 0.712-0.802) | **<2.2e-16** |
| Males with X*Y females<sup>1</sup> | **Y: 0.359** (95% CI: 0.311-0.409) | **3.28e-08** |
| XX* females | **X: 0.465** (95% CI: 0.413-0.517) | 0.19 |
| X*Y females<sup>1</sup> | **X\*: 0.467** (95% CI: 0.434-0.511) | 0.14 |

<sup>1</sup>As X\*Y females produce lethal YY embryos, to determine the transmission ratio of sex chromosomes of males and females involved in these crosses, we compared only the proportion of XX\* vs. X\*Y offspring (248 vs. 139) and XX\* vs. XY offspring (248 vs. 283) respectively.

### Impact of male sex chromosome drive on the stability of the system

To better understand the relation between the evolution and maintenance of the X* in *Mus minutoides* and the sex chromosome drive described in this study, we modeled the evolutionary dynamics of this system with a set of population genetics models. Our first aim was to determine the conditions allowing the maintenance of the X*, in the light of our new results. Based on standard stability analysis procedure (40) (see *Appendix A* in Supp. text), we show that the system is stable as long as the fertility of sex-reversed females (*w*_*X*\**Y*_, in numberof zygotes produced), exceeds a critical threshold (*w*_*crit*_), which value depends on the transmission ratio of males’ Y chromosome in crosses with XX and XX* females (*k*) and with X*Y females (*k\**):

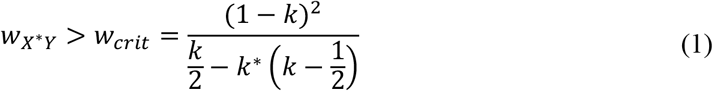

First off, equation (1) reveals that the X* is more likely to be maintained for greater values of *w*_*X*\**Y*_. In the absence of sex chromosome drive (*k* = *k\** = 0.5): the X* can be maintained as long as X*Y females have a fitness advantage over XX and XX* females (*w*_*X*\**Y*_ > *w*_*crit*_ = 1). This result replicates Bengtsson’s finding (32), who demonstrated that the loss of YY embryos does not select against the X*: with 1:1 segregation and *w*_*X*\**Y*_ = 1, X*Y females produce as many X*-bearing offspring as XX females produce daughters, even though YY are lost. As 2/3 of the offspring of sex-reversed females inherit the X*, even the slightest compensation for the loss (*w*_*X*\**Y*_ > 1), whether it is “automatic” (decreased competition between surviving embryos) or evolved (*e*.*g*. increased ovulation rate of X*Y females), will provide a selective advantage to the X*. In the presence of drive, the value of *w*_*crit*_ decreases when *k* increases (fig. 1): the more males transmit their Y in crosses with XX and XX* females, the easier is the X* maintained. In particular, with Y-drive (*k > 0*.*5*), the X* can be maintained despite lower relative fertility of X*Y females (*w*_*X*\**Y*_ < 1). The reason is two-fold: the X* is advantaged over the X because (i) it resists the drive (X*Y females transmit their X* and Y equally), and (ii) it allows to produce more females, the rarer sex in a context of Y-drive (see *Appendix A* in supp. text). The impact of the transmission ratio of male sex chromosome in crosses with X*Y females (*k\**) is less crucial, and varies depending on the value of *k* (fig. 1): with *k > 0*.*5* (Y-drive), if *w*_*X*\**Y*_ > 1, the system is stable regardless of *k\**, and if *w*_*X*\**Y*_ < 1, the stability is facilitated by small values of *k\**, which result in a decrease in production of the less fit X*Y females (see table 1). With *k < 0*.*5* (X-drive), the X* can only be maintained if *w*_*X*\**Y*_ > 1, and stability is favored by high values of *k\**, which increase the production of the fitter X*Y females.

**Figure 1.**
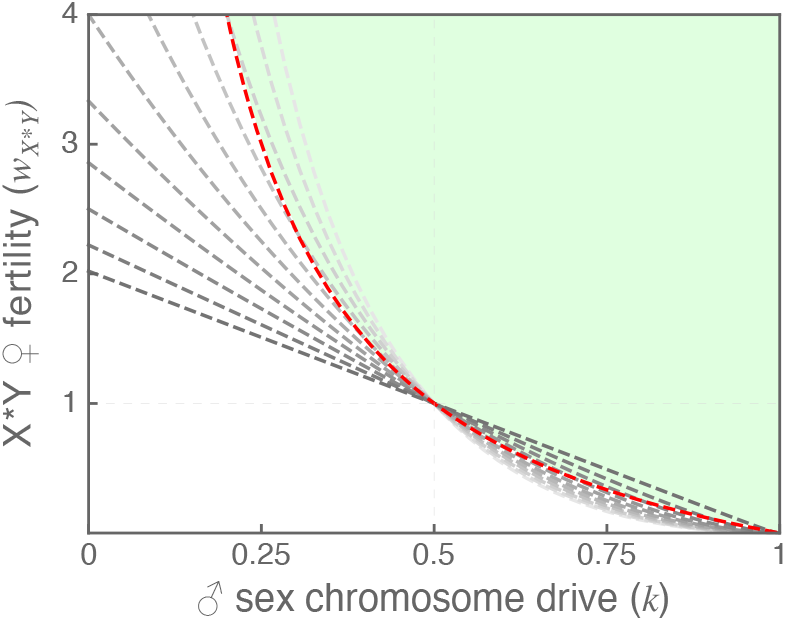
Stability of the polygenic sex determination system. The dashed lines represent *w*_*crit*_, the critical fertility value for X*Y female above which the X* chromosome can be maintained (see equation (1)), as a function of *k*, the transmission ratio of male sex chromosomes in crosses with XX and XX* females. The different curves show *w*_*crit*_ for different values of *k\** (transmission ratio of male sex chromosomes in crosses with X*Y females), ranging from 0 to 1 (gray to black scale, 0.1 increment). The red curve is *w*_*crit*_ for *k=k\** (unconditional drive), the green area depicts the parameter space in which the X* is maintained.

Using values of *k* and *k\** measured empirically (rounded to 0.8 and 0.36) into equation (1) provides an estimation of the minimum fertility of X*Y females allowing for maintenance of the polygenic system in the African pygmy mouse: *w*_*X*\**Y*_> 0.137 *i*.*e*., less than 1/7th of the fertility of XX and XX* females. No estimation of *w*_*X*\**Y*_ is available from wild populations, but knowing that X*Y females have a greater reproductive output in laboratory conditions (26), it is safe to assume that the X* is stable in natural conditions in this species, at least in part thanks to sex chromosome drive.

Our model also allows us to estimate the equilibrium frequencies of males and of the three types of females (Fig. S1; see *Appendix A* in supp. text). With increasing values of *w*_*X*\**Y*_, the frequencies of XX* and X*Y females increase whereas that of XX females and males decrease. With *k*=0.8 and *k\**=0.36, the model predicts that population sex ratio would be even for *w*_*X*\**Y*_=0.57, and slightly female-biased for greater values of *w*_*X*\**Y*_. It also predicts that XX females should be rare, less than 7% for *w*_*X*\**Y*_ > 1. This prediction is in line with field observations: only one out of 20 females captured in Caledon Nature Reserve (where the founder individuals of our laboratory colony were collected) had a XX genotype (27, and additional unpublished data).

### Paths to the evolution of a polygenic sex determination system with conditional sex chromosome drive

In this part, we evaluate the plausibility of different scenarios to explain the transition from a standard XX/XY sex determination system with no sex chromosome drive to a polygenic sex determination system with conditional drive of male sex chromosomes, as found in the African pygmy mouse. These scenarios consist of a sequence of events, with several steps (mutations) necessary to achieve the full transition, as shown on fig. 2. On the left side of fig. 2 are two scenarios in which the X* appears following the establishment of Y-drive (step a1). This X*, in addition to its feminizing effect, either has a direct effect on the transmission ratio of male sex chromosomes in crosses with X*Y females (step a2), or not (step a2’). In the latter case, its spread would have to be followed by the invasion of a drive modifier, affecting the transmission of male sex chromosomes in crosses with X*Y females specifically (step 3). On the right side of fig. 2 are two scenarios in which the X* emerges in a XX/XY population with no pre-existent sex chromosome drive. Conditional drive evolves once the feminizing chromosome is established, either in a single step (step b2), or in two steps: a first “non-conditional” driver invades, which has the same effect in all crosses (step b2’), followed by a drive modifier (step 3). For each step in these scenarios, we derived the conditions allowing for the spread (and fixation when relevant) of a mutant allele leading from one state to the next, using standard equilibria and stability analyses (40). All models and results are provided in *Appendix* B in supp. text, and are discussed in more intuitive terms in the following.

**Figure 2.**
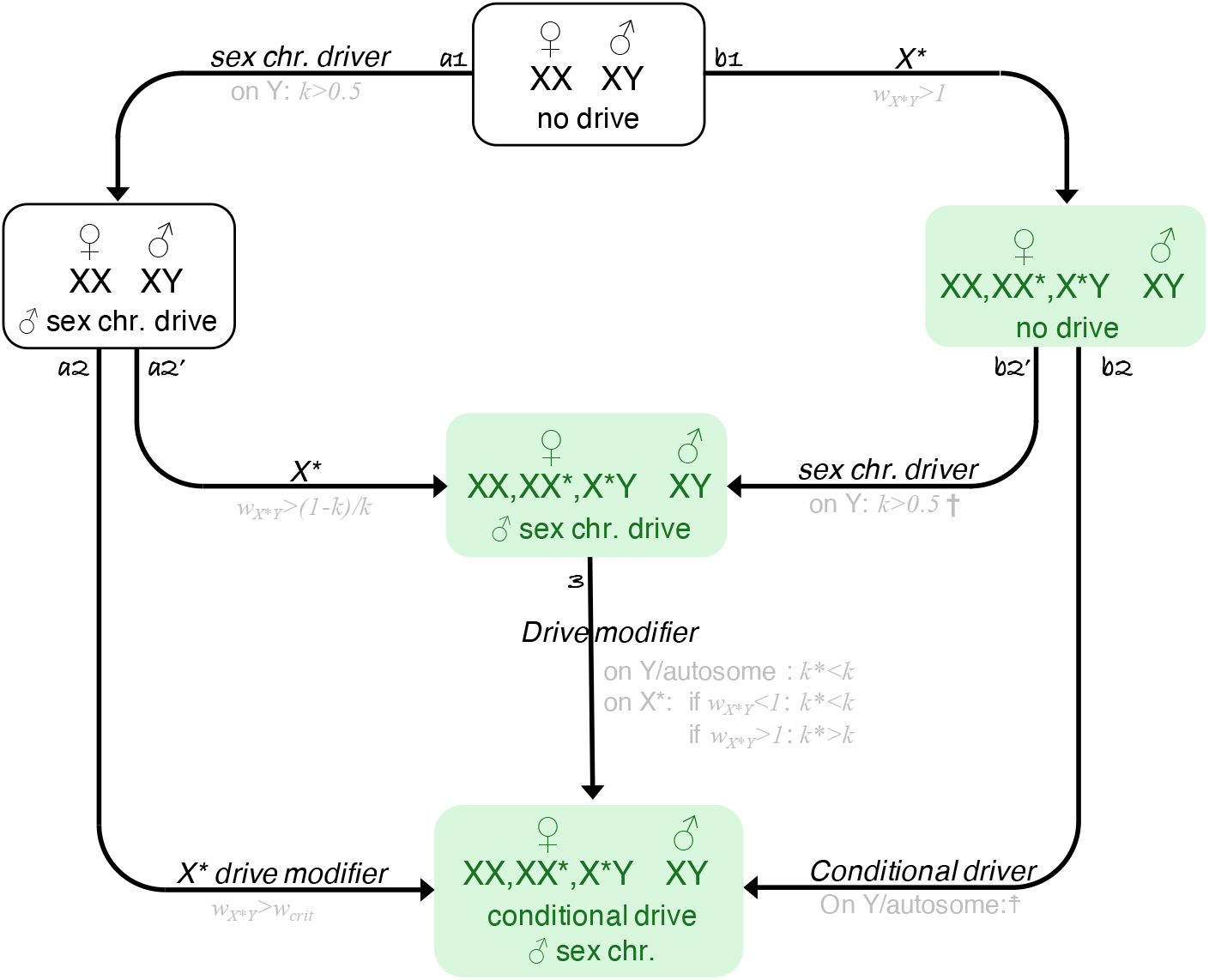
Evolutionary scenarios for the transition from standard male heterogametic system with no sex chromosome drive to a polygenic sex determination system with conditional drive of male sex chromosomes. Each square represents a state along the transition, green shaded squares are states at which the X* is present. Arrows indicate the type of mutation involved in the step between two state, along with the genomic compartment on which the mutant has to be for a successful invasion as well as the condition allowing invasion and fixation when relevant. †: if autosomal, a sex chromosome driver will increase in frequency when rare for *k>*0.5, but will stay at low frequency across most of *k, k\** and *w*_*X*Y*_ parameter space considered in the models (see fig. S2). ☨: a Y-linked or autosomal conditional driver will spread when rare for *k>*0.5, but might stay at an intermediate frequency, depending on the values of *k, k\** and *w*_*X*Y*_ (see fig. S3).

In brief, we show that the four speculative evolutionary paths are theoretically possible; for each step, the genomic compartment(s) on which the mutant considered can invade (and go to fixation when relevant), and conditions under which they can, are shown on fig. 2. In agreement with models by Kozielska et al. (36), we found that the spread of the feminizing X* is facilitated by the presence of a Y-drive in a standard XX/XY system: in the absence of drive, the X* can only spread if X*Y females have a greater fertility than the others (*w*_*X*\**Y*_ > 1, step b1), while if a Y-drive pre-exists, an emergent X* can spread despite a reduced fertility of X*Y females. If the X*, in addition to its feminizing effect, modifies male sex chromosome drive in XY x X*Y crosses (step a2), the condition for spread is *w*_*X*\**Y*_ > *w*_*crit*_ (the stability condition discussed in the previous section, eq. (1)), with a critical fertility always smaller than 1 if *k>0*.*5*. If the X* does not affect male sex chromosome drive (step a2’), it will invade the population for:

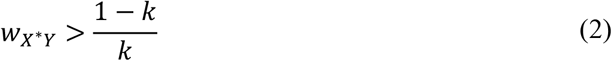

a simplification of eq. (1), with *k = k\**, also smaller than 1 if *k>0*.*5* (fig. 1).

In contrast, the spread of Y-linked sex chromosome drivers is not facilitated by the presence of an X*: a driving-Y will replace a non-driving Y only if it favors its own transmission (*k*>0.5, step b2’), the same condition found in populations with standard male heterogametic sex determination (step a1; (30, 41)). Nevertheless, as shown previously by Bull and Bulmer (34, 42), autosomal alleles favoring the transmission of males’ Y chromosome can also be selected in the presence of the X*. The reason invoked is that such driving alleles allow males to produce more sons on average, the rarer sex in that context. However, they never explored the fate of these alleles: can they spread to fixation? With our models, we replicated Bull and Bulmer’s results: a dominant autosomal driver of male sex chromosomes would indeed invade when rare, under the same conditions as a Y-linked driver (*k*>0.5). Using deterministic simulations, we show that the mutant allele would likely stay at a very low frequency, unless the strength of drive is very mild (fig. S2). For instance, for *k=*0.8, the maximum equilibrium frequency of the driving allele is less than 0.05, making it unlikely to go to fixation. The reason is that such an autosomal allele is under conflicting pressures: when it is rare, males produce more sons on average, via breeding with XX and XX* females. But as it increases in frequency, so do X*Y females, and soon, the advantage of giving birth to more sons in crosses with the first two types of females is outweighed by the cost of producing more daughters in crosses with X*Y females.

As mentioned earlier, conditional drive could have evolved in a single step (step b2), or in two steps: first with the spread of a first “non-conditional” driver, followed by the evolution of a drive modifier affecting male sex chromosome transmission specifically in XY x X*Y crosses (step 3). Mechanistically, there are different ways in which a conditional drive can evolve, our goal was not to be exhaustive and we considered the following scenario. We assumed that the mutant allele has distinct effects in males and X*Y females: in males, it causes a sex chromosome drive of strength *k* (e.g. through meiotic drive), in X*Y females mating with “driving males”, drive becomes *k\** (e.g. through a cryptic choice mechanism interfering with the fertilizing ability of sperm cells carrying the mutant allele). Interestingly, we show that different genomic compartments could carry the mutation(s) causing conditional drive. If it evolves in one step (step b2), the driver could successfully go to fixation if carried by the Y or an autosome. A rare conditional driver will increase in frequency for *k*>0.5, fixation is possible in both cases, though the conditions for the fixation of an autosomal sex chromosome driver are more restrictive (fig. S3). An autosomal driver can only fix if it favors the X in crosses with X*Y females (*k*<*0.5), as it allows to produce more of the rarer sex (males) and reduces proportion of YY embryos, which are both beneficial from an autosomal point of view. If it evolves in two steps, the secondary drive modifier could evolve on any of the three nuclear compartments found in X*Y females (step 3): an autosome, the Y or the X*. If the drive modifier is autosomal or Y-linked, it goes to fixation assuming it decreases Y-drive (*k*^*^ < *k*), as this reduces the proportion YY embryos produced, and increases the proportion of males. If X*-linked, it will fix if *k*^*^ > *k* when *w*_*X*\**Y*_ > 1, as the mutant X* gains a fitness advantage by producing more of the fitter X*Y females, and assuming *k*^*^ < *k* when *w*_*X*\**Y*_ < 1, for opposite reasons.

### The long term persistence of the polygenic sex determination system

As long as the system is ecologically stable, the X* is protected against loss, and so are the X and Y chromosomes (they are both essential to produce males). As explained by Maynard-Smith and Stenseth (43), to be stable in an evolutionary sense, the system has to be able to resist the introduction of genetic modifiers suppressing the feminizing activity of the X*. Such suppressors could arise on the Y chromosome or an autosome, and in their presence, X*Y individuals would develop as males, which in turn would lead to the production of X*X* females. Ultimately, the spread of a suppressor could theoretically drive the system to revert to standard male heterogamety, with either X*X* females and X*Y males or XX females and XY males, following the loss of either the X or X* chromosome (fig. 3). We used standard stability analyses to determine the conditions under which the X-X*-Y system is evolutionary stable in the presence of conditional male sex chromosome drive, and for conditions under which the system is unstable, we simulated how the spread of a suppressor influences sex determination. Results are fully detailed and discussed in *Appendix C* in supp. text, and here we describe the main results.

**Figure 3.**
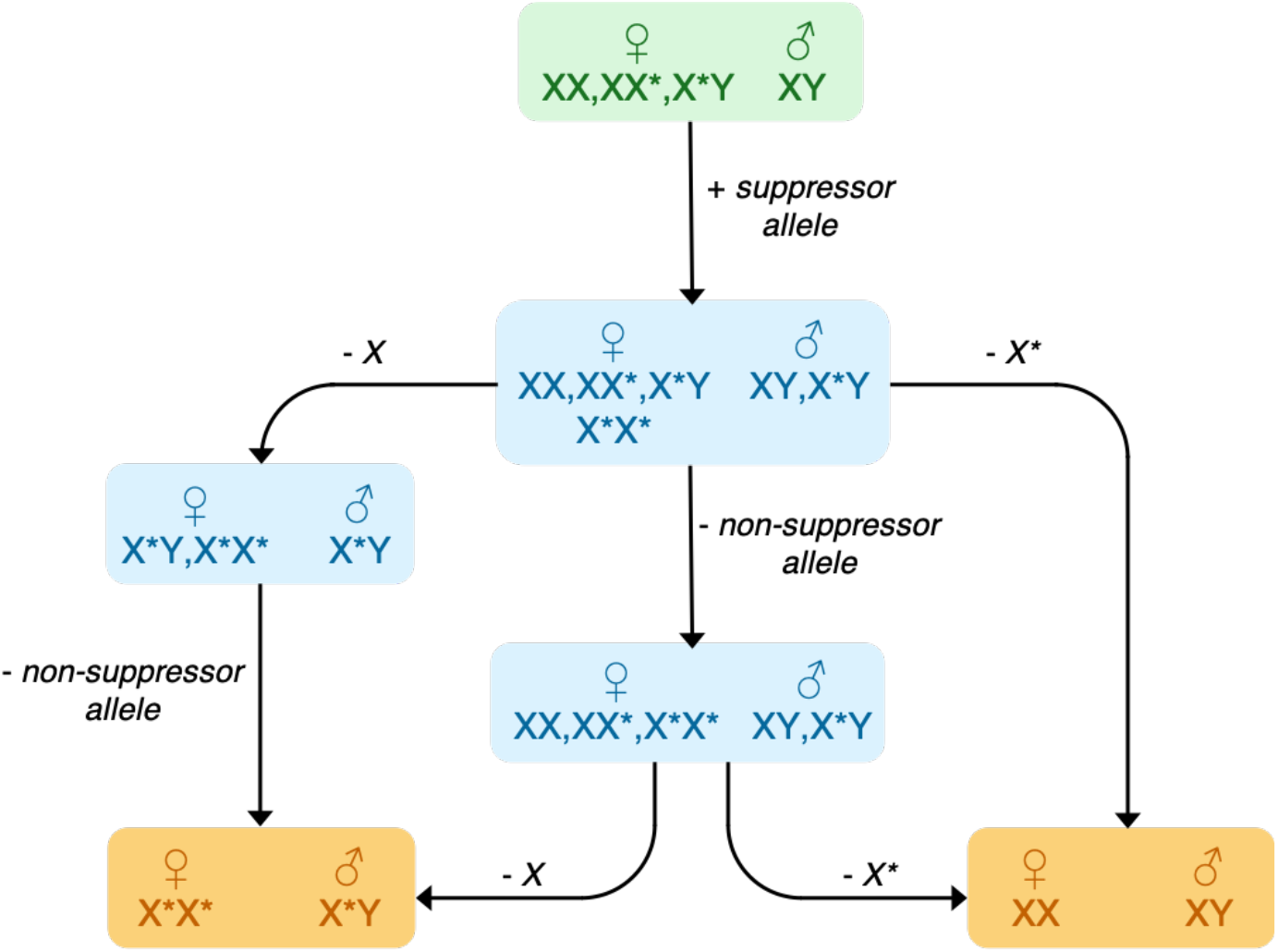
Paths leading to heterogamety following the spread of a suppressor of the feminizing activity of the X*. Worth for both a Y-linked or autosomal suppressor. Each square represents a putative equilibrium state, and color indicates the condition of sex determination: green: polygenic sex determination currently found in the African pygmy mouse, blue: “alternative” polygenic sex determination, orange: male heterogamety. Arrows leading from one state to the next indicate which allele is lost (-) or gained (+) between two states. Following the spread of suppressor, several “alternative” stable polygenic state, at which more than two sexual genotypes persist in at least one of the two sexes, could be reached: first, if polymorphism is maintained at the suppressor locus in the presence of X, X* and Y sex chromosomes, four types of females (XX, XX*, X*Y and X*X*) and two type of males (XY and X*Y) co-exist. Second, if the suppressor allele goes to fixation, but the three sex chromosomes are maintained at equilibrium, a system with XX, XX* and X*X* females and XY and X*Y males can be established. Finally, the loss of the X chromosome would result in a polygenic system with X*Y and X*X* females and X*Y males. Reaching a X*X*/X*Y male heterogametic system requires the loss of the X and fixation of suppressor allele, and reaching a XX/XY system is achieved following the loss of the X* (even if the suppressor locus remains polymorphic).

Conditions that allow the spread of a rare suppressor are similar whether it is Y-linked or autosomal (fig S4-5): a low fertility of X*Y females (hereafter *w*_*X*\**Y♀*_) and high fertility of X*Y males (*w*_*X*\**Y♂*_) (and X*X* females (*w*_*X*\**X*\*_) if it is autosomal, fig S6) will tend to favor its invasion. Nevertheless, invasion of a suppressor is easier (possible across a greater range of parameter values) if it is Y-linked. Concerning the impact of male sex chromosome drive: the strength of drive in crosses with XX and XX* females (*k*) only has a minor impact on evolutionary stability, as opposed to the strength of drive in crosses with X*Y females (*k\**): the spread of suppressors is hindered by low values of *k\** and favored by high values (fig S4-5). In other words, an unconditional Y-drive of male sex chromosomes reduces evolutionary stability (as opposed to no sex chromosome drive), while a conditional drive such as the one found in the African pygmy mouse (*k*=0.8 and *k\**=0.36), favours stability (fig 4).

**Figure 4.**
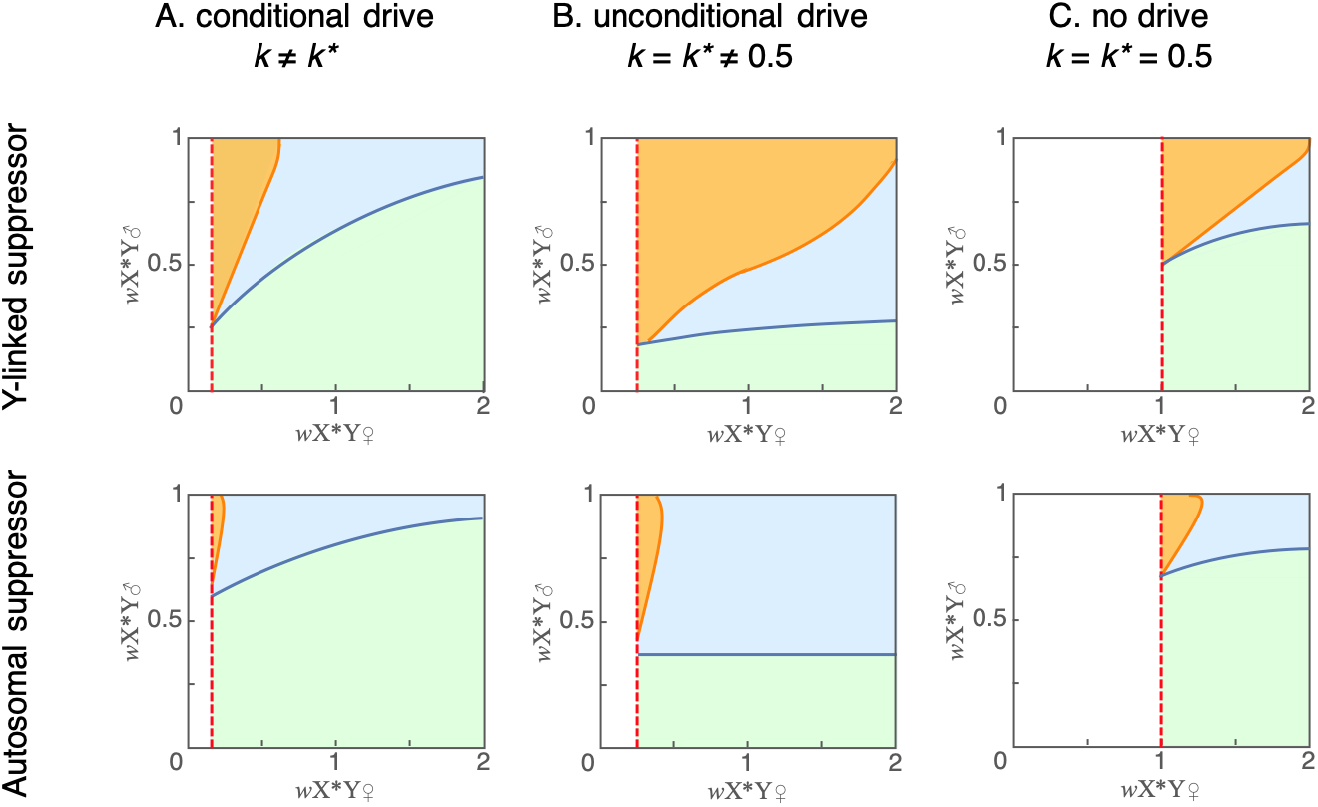
Evolutionary outcomes in the presence of a suppressor of the feminizing activity of the X*, as a function of the fertility of X*Y females and X*Y males (fertility of X*X* females fixed to one). The suppressor is either Y-linked (top row) or autosomal (bottom row), with either (A) conditional drive of male sex chromosomes (*k*=0.8 and *k\**=0.36), (B) unconditional drive of male sex chromosomes (*k*=*k\**=0.8) and (C) no drive of sex chromosomes (*k*=*k\**=0.5). We delineated three regions in the parameter space that correspond to qualitatively different evolutionary outcomes of the introduction of a suppressor (see fig. 3). In the green areas, the system is stable against its invasion. In the blue areas, a rare suppressor can spread but does not cause a return to a standard male heterogametic sex determination. In the orange areas, a suppressor will spread and cause a return to male heterogamety (XX/XY or X*X*/X*Y). The boundaries shown are numerically predicted representations based on the analytical models and deterministic simulations presented in Appendix C. The red dashed line shows the threshold for maintenance of the X*, which depends only on the relative fertiliy of X*Y females (see equation 1).

If the conditions to resist spread of a suppressor are not met, our models show that the system can either reach a stable alternative polymorphic state, or revert back to male heterogamety (fig. 3). Alike the conditions for invasion, a suppressor is more likely to cause a return to male heterogamety if *w*_*X*\**Y♀*_ is low and *w*_*X*\**Y♂*_ high, and if it is Y-linked. With a conditional drive of male sex chromosomes (*k*=0.8 and *k\**=0.36), a return to male heterogamety would only be possible if X*Y females had a very poor fertility compared to other females: around *w*_*X*\**Y♀*_ = 0.5 for a Y suppressor and *w*_*X*\**Y♀*_ = 0.25 for an autosomal suppressor (fig. 4A). As X*Y females in the pygmy mouse have a higher reproductive output than XX and XX* females in laboratory conditions (26), it is unlikely that their fertility in the wild would be so low. This means that if ever a suppressor of X* activity emerged, it might be able to spread to an intermediate frequency, but it would not cause a loss of the polygenic sex determination system. Interestingly, if the drive were unconditional (*k=k*=*0.8, fig. 4B) or inexistent (*k=k*=*0.5, fig. 4C), the system would be more vulnerable to suppressors: the regions of the parameter space in which (i) suppressors can invade and (ii) a standard male heterogamety is restored are larger (*i*.*e*., invasion of the suppressor and loss of the X or X* would be possible for relatively higher fertilities of X*Y females and lower fertilities of X*Y males). These results suggest that in African pygmy mouse, the polygenic sex determination system is protected by the conditional drive of male sex chromosomes: it is less likely to be compromised by a suppressor.

## Discussion

There is growing evidence that selfish genetic elements are important drivers of evolutionary innovation (29, 44–50). Sex chromosome drive, and especially Y-drive, is nevertheless rare, supposedly because of its impact on population sex ratio, which triggers the evolution of suppressors (27, 29, 51). In this paper, we describe a remarkable system of male sex chromosome drive in the African pygmy mouse, which direction (X-drive or Y-drive) and strength depend on the genotype of the male’s sexual partner.

By combining our empirical findings with mathematical modeling, we demonstrate that the sex chromosome drive in the African pygmy mouse helps maintaining the feminizing X* chromosome, and that the conditional nature of the drive is crucial in limiting the spread of suppressors of X* activity: this unusual sex determination system is “locked in” thanks to the biased transmission of male sex chromosomes. Our models further confirm that the X* could have evolved in response to a selfish genetic element biasing the transmission of male sex chromosomes (*i*.*e*., as a mechanism for resistance to drive), in agreement with previous theoretical work (36, 37). Nevertheless, the X* could have emerged in the absence of sex chromosome drive (which would thus have evolved secondarily), provided that X*Y females had a greater fertility than XX and XX* ones at its inception. Although X*Y females were shown to have a higher reproductive output (26), it is likely that they originally had a poor fitness (XY females in mammals, including the laboratory mouse, tend to have poor fertility if not completely sterile (13, 15)). They could have acquired their fitness advantage subsequently, thanks to a gradual accumulation of female-beneficial genes and alleles on the non-recombining region of the X*, as could be expected considering the canonical model of sex chromosomes evolution (52), and/or on the Y (38). It is also possible that neither sex chromosome drive nor a greater fecundity of X*Y females was the initial trigger for the spread of the X*: some theoretical models show that interdemic selection (53) or strong inbreeding (54) can favor the spread of sex-reversal genes. For now, too little is known about the ecology of the African pygmy mouse (55) to declare if these hypotheses are relevant or not. To go further, it would be valuable to study other pygmy mice populations. X*Y females in this species have been found from Southern up to Western Africa, and the proportion of X*Y females seems to vary across localities (56), suggesting sex chromosome drive might differ from one population to another. Comparing the transmission ratio of male sex chromosomes and breeding success of X*Y females in different regions could help further disentangle the cause(s) of the evolution of the X* in this species.

As mentioned in the introduction, X*Y females are found in several other mammalian species, and sex chromosome drive is found in one shape or another in all of them. A conditional male sex chromosome drive system similar to the one of the African pygmy mouse seems to exist in the mandarin vole *L. mandarinus* (11). What is remarkable is that the direction and strength of drive appears to fit with our observations in the African pymgy mouse: the Y was estimated to be transmitted to close to 80% of offspring in crosses with XX and XX* females, while it is transmitted at a rate of around 10% in crosses with X*Y females (note that estimations are based on a limited number of genotyped offspring, so they remain to be confirmed). The conclusions drawn from our mathematical models for *M. minutoides* are therefore also largely valid for *L. mandarinus*. In those two species, and all other species with X*Y females, whether the evolution of sex chromosome drive predates, follows (or even coincides with) the emergence of the X* remains to be established, but the recurrent co-occurrence of the two is clearly puzzling. In standard heterogametic sex determination systems, sex chromosomes are hot-spots for genomic conflicts because of their peculiar transmission patterns. The presence of a third sex chromosome increases the number of genomic compartments that segregate independently and can engage in genomic conflicts, so polygenic sex determination systems might be more prone to the emergence and spread of sex-chromosome drivers. Our models and others (32, 57) actually suggest that these systems are more tolerant towards the invasion of certain types of sex chromosome drivers. For instance, in standard XX/XY and ZZ/ZW systems, sex-ratio selection tends to favours autosomal loci that insure a balanced transmission of sex chromosomes, as illustrated by the autosomal suppressors of sex chromosome drive found in several *Drosophila* species (49, 58, 59). In contrast, in systems such as the one found in the African pygmy mouse, because the X* turns certain genotypic males into females, and therefore biases sex ratio, it might not be in the best interest of autosomes that sex chromosomes are transmitted equally, as shown by the fact that a Y-chromosome driver carried by an autosome can be selected under certain circumstances (fig. 2, steps b2 and b2’). Furthermore, our models show that different genomic compartments could harbour mutations limiting the transmission of male’s Y chromosomes in crosses with X*Y females (fig. 2, step 3): autosomes, the X* and even the Y itself. For autosomes, in addition to increasing the proportion of males, this is also valuable as it reduces the probability of ending up in non-viable YY embryos. For the X*, it is only true if the fertility of X*Y females is lower than that of XX* females, as it increases its chances to be associated with an X chromosome. Finally, from the Y’s perspective, it is also advantageous to increase the transmission of the X at its own expense. Its net transmission ratio (number of viable embryos the Y chromosome ends up in) is unchanged, as X*Y females also pass down Y chromosomes, but contrary to males that transmit their Y exclusively to X*Y embryos, female pass it down to males only, providing a fitness advantage in a context of female-biased sex ratio. These findings illustrate that the interests of different genomic compartments that are usually in conflict over the transmission of sex chromosomes are modified in the presence of a third sex chromosome, and that these interests can even align in these specific cases.

The conditional nature of the drive in *Mus minutoides* also raises many questions regarding the underlying proximal mechanism(s). Assuming that the X* evolved in response to a Y-chromosome drive, one possibility in line with our analytical results is that in all males, a selfish element on the Y chromosome promotes its transmission through meiotic drive or by interfering with maturation of sperm cells harboring the X chromosome, making them dysfunctional (the most widespread mechanism for chromosome drive (60)). The X-drive specific to crosses with X*Y females could have evolved subsequently, and for instance, females could exercise a cryptic choice to favor fertilization by X-bearing sperms (*i*.*e*., by rendering the genital environment hostile to Y-bearing sperm), or selectively abort “unwanted” embryos (e.g. through maternal imprinting (11)). As the biased transmission of male sex chromosomes was identified based on genotyping pups at weaning, a profusion of mechanisms, ranging from meiotic drive to a differential mortality of embryos or pups bearing paternal X or Y chromosomes could be responsible for sex chromosome drive. Additional experiments are therefore necessary to pinpoint the exact mechanism(s) involved. Concerning genetic basis, most drive system seem to emerge from gene duplication events (49), and can involve massive gene amplification due to the concurrent evolution of drivers and suppressors of drive. For instance, one of the most comprehensively described drive system, found in the house mouse, involves an arms race between the sex-linked multicopy genes *Slx* and *Sly*, found in more than a hundred copies respectively on the X and Y chromosome (61, 62). Assessing the presence and copy number of these genes (and other post-meiotically expressed genes) on the three sex chromosomes of the African pygmy mouse might be a good way to begin investigating the genetic architecture underlying the conditional drive of in this species. Clearly, rodents with feminizing X chromosomes are remarkable on many levels. They appear to be particularly prone to the accumulation of sex chromosome drivers, as illustrated by the conditional drive of male sex chromosomes of the African pygmy mouse, described in this paper. *Mus minutoides* and the other mammals with unusual sex determination systems make excellent models to study genomic conflicts, and in particular the proximal mechanisms and genetic basis of sex chromosome drive, which are still poorly understood.

## Materials and Methods

### Sex ratio at weaning and sex chromosomes transmission ratios

In June 2010, a breeding colony of *Mus minutoides* was established from animals caught in Caledon Nature Reserve, South Africa (for full details see references (9, 26)). New couples were systematically formed after weaning, and breeding was closely monitored. Progeny sex ratio data was acquired during four consecutive years, from 27, 49 and 73 couples with respectively XX, XX* and X*Y females, for a total of 74, 130 and 194 litters. The number of males and females in each litter was determined at weaning, the two sexes being unambiguously told apart based on ano-genital distance and general external genitalia appearance (25). The sex chromosome complement of females was then assessed by PCR amplification of the Y-specific *Sry* gene and/or karyotyping; as previously described (9).

Sex ratio (defined here as the proportion of males) at weaning was assessed for the three types of crosses and compared to expected sex ratios under the hypothesis of Mendelian transmission (0.5 for XX females, 0.25 for the XX* and 0.33 for the X*Y) with binomial tests. The transmission ratio of X and Y chromosomes in males in each type of crosses, and of X and X*, and X* and Y, in respectively XX* and X*Y females was measured, and departures from the expected 50:50 transmission ratio were tested with binomial tests.

### Theoretical analyses

The mathematical models developed in this study were inspired by models developed to study sex determination in the lemmings *Myopus schisticolor* and *Dicrostonyx torquatus* (32, 34, 35, 43, 63–65), and adapted to fit *Mus minutoides’* distinctive features. Here we provide the outline of the model and describe the main procedures. All details can be found in supplementary material.

#### The model

The model is a standard population genetics model, which assumes an infinite diploid population with random mating and non-overlapping generations: a system of recurrence equations gives the frequencies of male and female genotypes at each generation, depending on their frequencies at the previous generation (see *Appendix A* in supp. text). Sex is determined by a single locus with three alleles (X, X^*^ and Y), the female determiner X* is dominant over the male determiner Y, which itself is dominant over the X. This results in the production of one type of males (XY; YY males are unviable) and three types of females (XX, XX^*^ and X*Y). In agreement with previous observations showing that XX and XX* females have the same reproductive output, different from that of X*Y females (26), the fertility of XX and XX* females is set to 1 in our model, and that of X*Y females is denoted as *w*_*X*Y*_. Fertility in our models controls the number of zygotes produced by a female, so the relative number of offspring that X*Y females actually give birth to is calculated by subtracting the relative number of YY zygotes produced from *w*_*X*Y*_. This is identical to the very first model built to study sex determination in the lemmings (32). Finally, to match the empirical results described in the present paper, the transmission of sex chromosomes is always random in females, and the ratio of Y chromosomes transmitted by males is conditional upon female genotype. The strength of distortion (proportion of male Y chromosomes transmitted to the progeny) is denoted *k* in crosses with XX or XX* females, and *k*^*^ in crosses with X*Y females.

#### Stability of the system

The aim of the stability analysis is to define the parameter space (for the three parameters *k, k*, w*_*X*Y*_) that allows the maintenance of the X* in the general model described above, provided that no genetic modifiers are introduced (ecological stability *sensu* Maynard-Smith & Stenseth (43)). Following standard equilibrium analyses described in Otto and Day 2007(40), we analyzed the eigensystem of the transition matrix associated with the system of recurrence equations described above. The population converges towards an equilibrium, which genetic composition, including presence or absence of the X*, is given by the eigenvector associated with the highest eigenvalue of the transition matrix (see *Appendix A* in supp. text for detailed analyses).

#### Evolutionary scenarios

Several scenarios, each composed of several steps, are analyzed to explore how a standard XX/XY sex determination system with Mendelian transmission of sex chromosomes can evolve into a polygenic sex determination system with conditional male sex chromosome drive, such as found in the African pygmy mouse (fig. 2). For each possible step, our aim is to define the conditions that allow going from one state to the next, *i*.*e*., that allow the mutant allele involved in this step to spread when rare. We use equilibria and stability analyses as described by Otto and Day (40). The exact procedure used depends on whether the mutant allele considered at a given step arises on the X*, Y or an autosome. If the mutant is born on the X*: the dynamical equations describing the system are linear functions of the variables (because all males have the same genotype), so the conditions for invasion of the mutant X* can be obtained directly through the study of the eigensystem of the transition matrix of the system at that step. If the mutant is Y-linked or autosomal, the model consists of a non-linear combination of multiple variables, because the mutant allele can be found in both males and females, resulting in multiple genotypes in both sexes. In this case, we first determine the conditions under which a rare mutant will increase in frequency (invasion conditions), by performing a stability analysis of the model at the equilibrium where the emergent mutant allele is absent. This requires studying the Jacobian matrix at the equilibrium point of interest: if its leading eigenvalue is greater than one, the equilibrium is unstable *i*.*e*., a rare mutant will increase in frequency when rare. However, a successful invasion does not necessarily imply the loss of the ancestral allele: a mutant allele can spread until it reaches an intermediate frequency, producing a stable polymorphism. As we are interested in the conditions that allow the mutant allele to replace the resident allele, we also establish the fixation conditions of the mutant allele, by performing a stability analysis of the model at the equilibrium where the resident allele is absent. If all eigenvalues of the Jacobian matrix at that equilibrium are smaller or equal to one, the equilibrium is stable *i*.*e*., a rare ancestral allele would decrease in frequency until it is lost and the mutant fixed in the population (see *Appendix B in supp. text* for detailed analyses).

#### Long term persistence of the system

In this part we define the conditions under which a suppressor of the feminizing action of the X* can evolve and cause the loss of the polygenic sex determination system. Such a suppressor can arise either on the Y or an autosome, and results in masculinization of X*Y individuals, which produce X*X* daughters when mated to females that carry an X*. This requires extending the general mathematical model by introducing in our models either a novel allele at the sex-determining locus if the suppressor is Y-linked, or a novel autosomal locus with two alleles: a wild-type “non-suppressor” allele and a dominant suppressor allele, which causes X*Y individuals to develop as males. As of this point, relative fertility of X*Y females is denoted *w*_*X*\**Y♀*_, and two new parameters are added to the models: (i) the fertility of X*Y males (*w*_*X*\**Y♂*_), which is relative to the fertility of XY males, and that we assume to be smaller or equal to one, as X*Y males might bear a cost for carrying the X*, (ii) the relative fertility of X*X* females (*w*_*X*\**X*\*_), which for simplicity, was either set to one (same fertility as XX and XX* females) or equal to *w*_*X*\**Y♂*_. These two cases make the most sense biologically considering the difference in reproductive success between XX and XX* females versus X*Y females. Case (i) is expected if the greater fertility of X*Y females stems from Y-linked genes or alleles, case (ii) if it stems from recessive X*-linked alleles. We decide to consider that males crossed with X*X* females see their Y chromosome transmitted with a ratio *k*, and that carrying the suppressor does not add any fertility cost, in order to avoid making our models too complex to analyze. We first tried to derive analytical conditions for stability against each type of suppressor (conditions under which the suppressors cannot invade), but the complexity of the models precluded obtaining analytical expressions for stability. We therefore derived them numerically, by replacing parameters by a wide range of numerical values. Then, we investigated the consequences on sex determination of the spread of the suppressor (see Fig. 4), by the mean of numerical deterministic simulations. (see *Appendix* C in supp. text for detailed analyses).

## Acknowledgements

This study was supported by the French National Research Agency (ANR grant “SEXYMUS”, No. 10-JCJC-1605), and the Del Duca Foundation from Institut de France (“Subvention scientifique 2012”). The experimental protocol was performed in accordance with European guidelines and with the approval of the Ethical Committee on animal care and use of France (N° CEEA-LR-12170). We thank the animal breeding facility of the University of Montpellier (CECEMA), M. Perriat-Sanguinet for his help in maintaining the breeding colony, and Sally Otto for valuable discussions.

