## Supplementary information for "Multiple Sex Chromosome Drivers in a Mammal with Three Sex Chromosomes"

### Supplementary information for:
### Multiple Sex Chromosome Drivers in a Mammals with Three Sex Chromosomes

Paul A. Saunders, Julie Perez, Ophélie Ronce and Frédéric Veyrunes

Corresponding author: Paul A. Saunders


This document contains supplementary figures cited in the main text (figures S1-S6) as well as three appendices (A, B and C) with all the details of the mathematical models. These appendices also cite figures S1-S6, and contain additional illustrative figures and tables, not cited in the main text, that are called using the prefixes A, B or C (*e.g.*, figure B1, table B1).

### supplementary figures

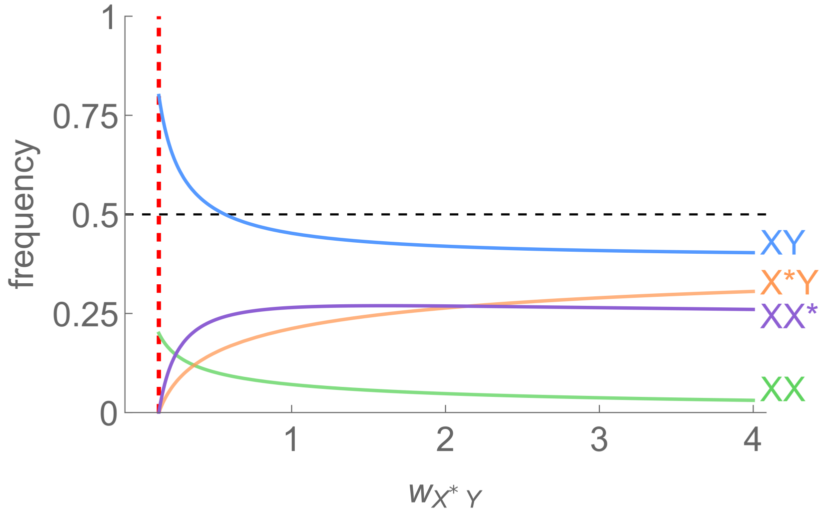

**Figure S1. Equilibrium frequencies of males and of the three types of females as a function of $w_{\mathbf{X^*Y}}$.** The values of $k$ and $k*$ were set to 0.8 and 0.36 to match empirical observations. The red dashed line shows the threshold for maintenance of the X* ($w_{X^*Y} = 0.137$). Frequencies were derived from models presented in Appendix A in supp. text.

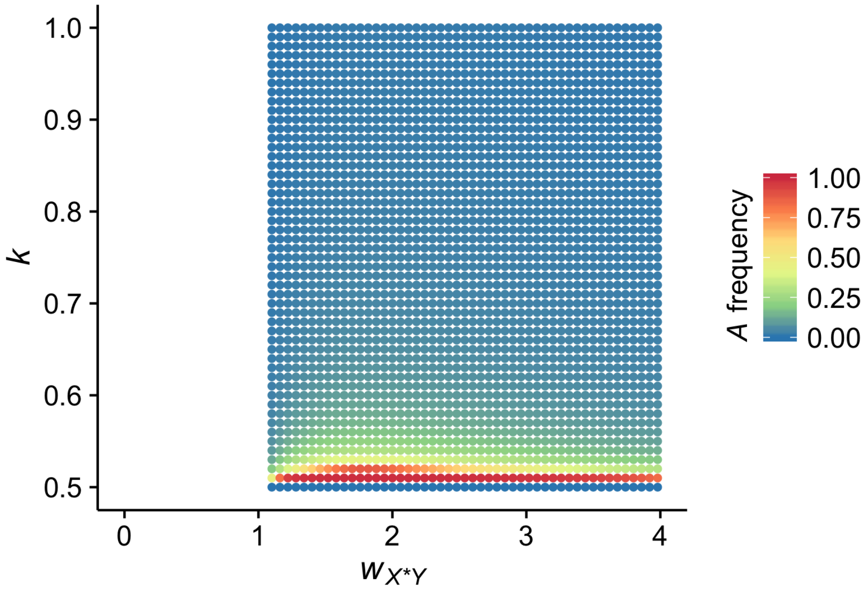

**Figure S2. Fate of a rare autosomal sex chromosome driver $\boldsymbol{A}$ in a XX,XX*,X*Y/XY system with no sex chromosome drive.** Each dot shows the outcome of a single numerical deterministic simulation with fixed values of $k$ and $w_{X^*Y}$. Simulations were started with a frequency of $A$ set to $10^{-3}$, and ran for 500,000 generations or until the change in frequency of $A$ was $< 10^{-20}$. Color indicates the frequency of the mutant $A$ at the end of the simulation. Maximum equilibrium frequency reached for $A$ is 0.987, for $k = 0.51$ and $w_{X^*Y} = 1.82$ (see Appendix B in supp. text for further details).

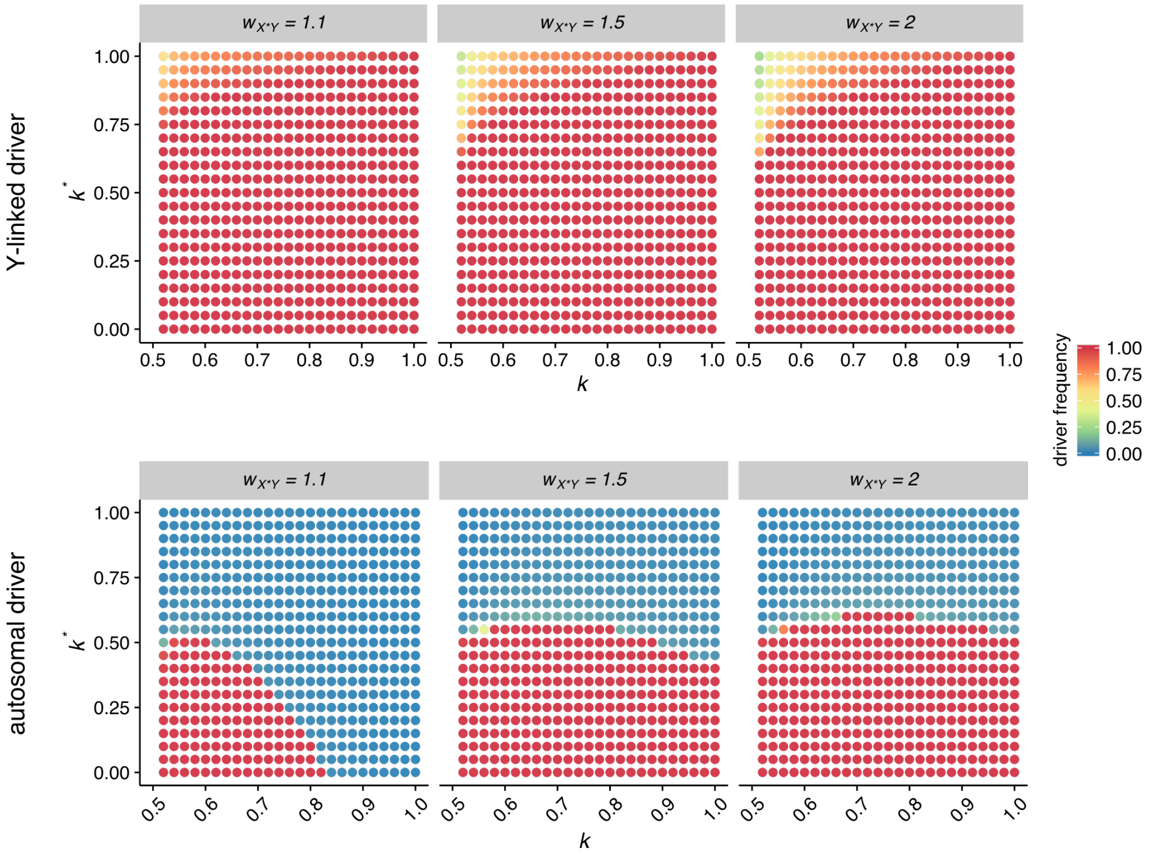

**Figure S3. Fate of a rare conditional sex chromosome driver in a XX,XX*,X*Y/XY system with no sex chromosome drive.** The driver is either Y-linked (top row) or autosomal (bottom row). Each dot shows the outcome of a single numerical deterministic simulation with fixed values of $k$ and $k^*$ and $w_{X^*Y}$. Simulations were started with a frequency of the driving allele set to $10^{-3}$, and ran for 20,000 generations or until the change in frequency of the driving allele was $< 10^{-20}$. Color indicates the frequency of the driving allele at the end of the simulation. (see Appendix B in supp. text for further details).

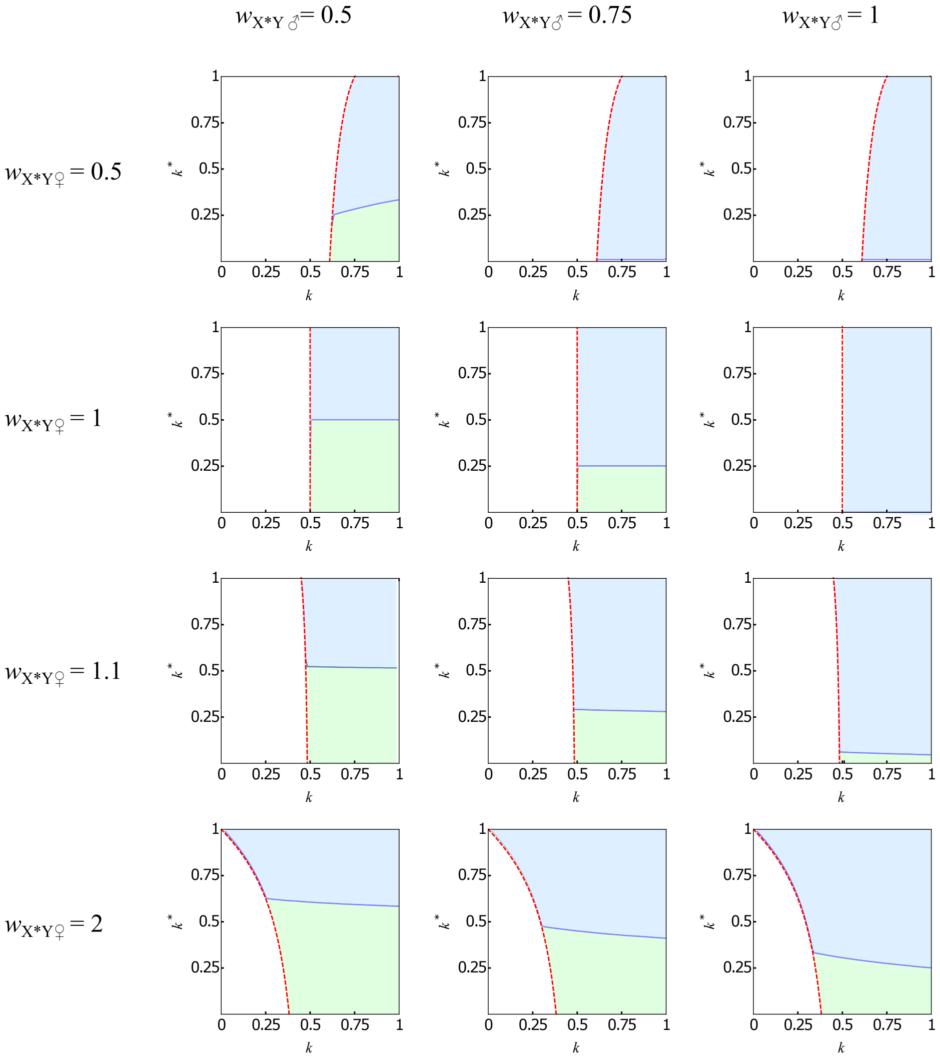

**Figure S4. Conditions for invasion of a Y-linked suppressor of the feminizing activity of the X*,** as a function of $k$ and $k^*$, for different set of values of $w_{X^*Y♀}$ and $w_{X^*Y♂}$. The red dashed line shows the threshold for maintenance of the X* (see Appendix A in supp. text). Green area: parameter space in which the suppressor Y cannot invade (stable system), blue area: parameter space in which it can invade (see Appendix C in supp. text for details).

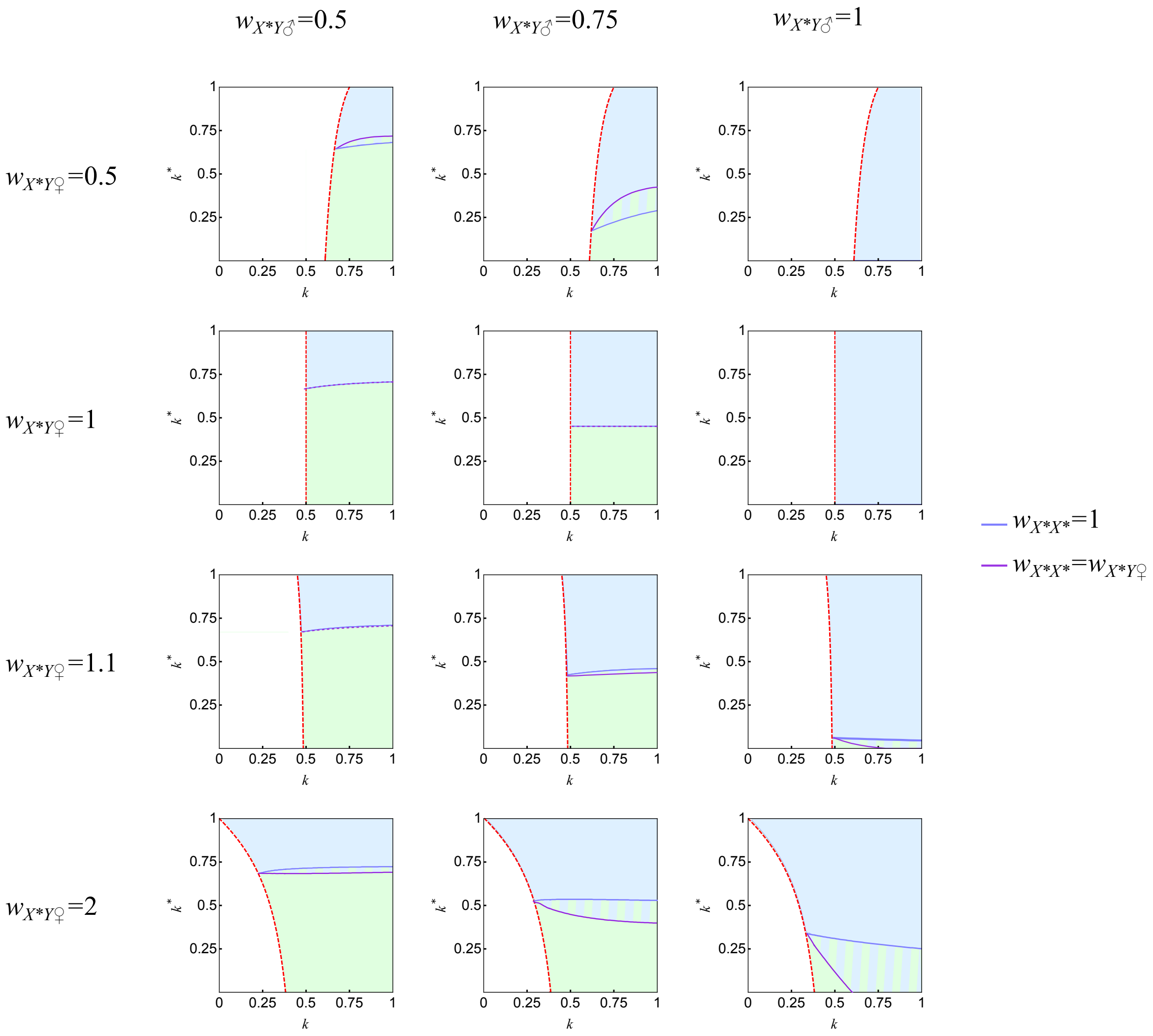

**Figure S5. Conditions for invasion of an autosomal suppressor of the feminizing activity of the X*,** as a function of $k$ and $k^*$, for different set of values of $w_{X^*Y♀}$ and $w_{X^*Y♂}$. The red dashed line shows the threshold for maintenance of the X* (see Appendix A in supp. text). Green area: parameter space in which the suppressor Y cannot invade (stable system), blue area: parameter space in which it can invade. (see Appendix C in supp. text for details).

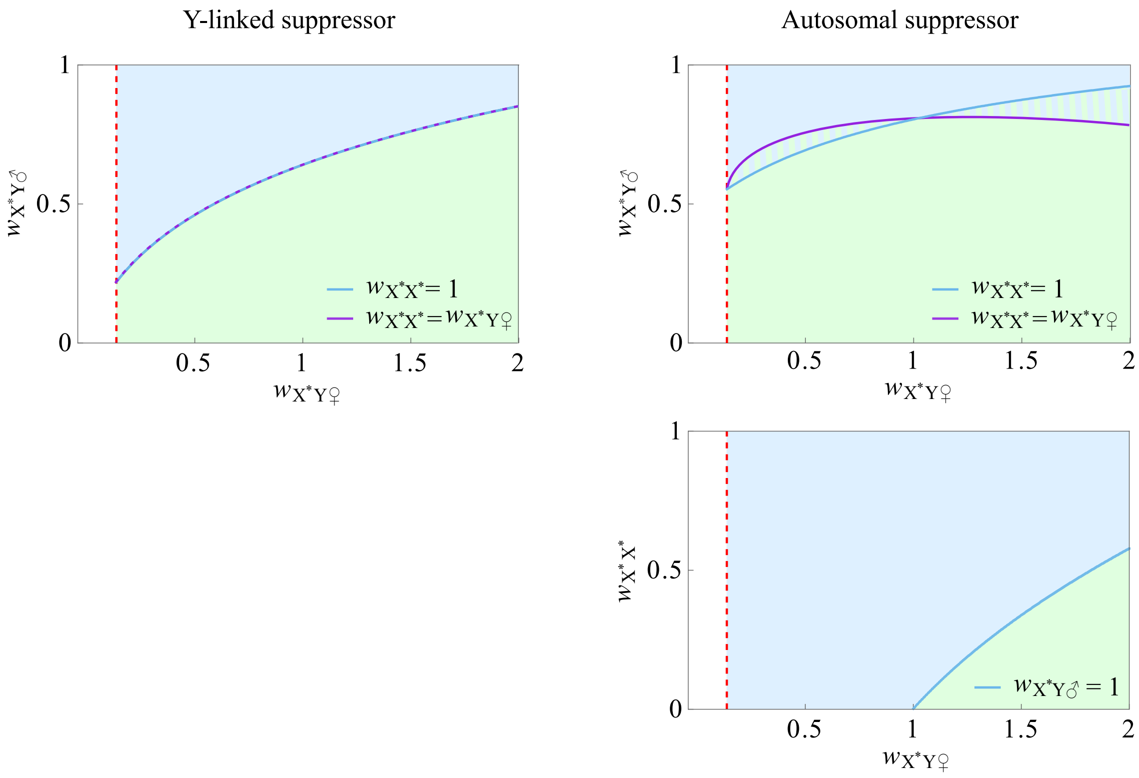

**Figure S6. Conditions for invasion of a rare suppressor of the feminizing activity of the X***, as a function of X*Y male fertility ($w_{X^*Y♂}$) and female X*Y and X*X* fertility ($w_{X^*Y♂}$, ($w_{X^*X^*}$)), with $k = 0.8$ and $k^* = 0.36$. The red dashed line shows the threshold for maintenance of the X*, which depends only on the relative fertility of X*Y females (see Appendix A in supp. text). The green area is the parameter space in which the suppressor cannot invade (stable system), the blue area in which it can invade (see Appendix C in supp. text for details).

### Appendix A - Stability analysis

This stability analysis aims at determining the conditions under which the polygenic sex determination is stable, *i.e.*, the conditions allowing the maintenance of the feminizing chromosome X*.

The model assumes three types of females: XX, XX* and X*Y in numbers $f_i$ ($f_1$, $f_2$, $f_3$) and one type of males (XY) in number $m_j$ ($m_1$). Females transmit their sex chromosomes in a Mendelian fashion, and the transmission ratio of male sex chromosomes depends on female genotype: the ratio of Y chromosomes transmitted is worth $k$ in crosses with XX and XX* females, and $k^*$ with X*Y females. We define fertility as the number of zygotes a female produces, assuming all but YY zygotes survive to adulthood and participate in producing the next generation. XX and XX* females are assumed to have the same fertility ($w_{\mathrm{XX}} = w_{\mathrm{XX^*}} = 1$) in line with empirical observations made in the African pygmy mouse. Fertility in X*Y females differs and is noted $w_{\mathrm{X^*Y}}$. As X*Y females produce a fraction $\frac{k^*}{2}$ of non-viable YY zygotes, we assume they produce $w_{\mathrm{X^*Y}}(1 - \frac{k^*}{2})$ viable offspring.

The dynamics of the number of females is given by the recursions:

$$\begin{bmatrix} f_1 \\ f_2 \\ f_3 \end{bmatrix}_{(t+1)} = M \begin{bmatrix} f_1 \\ f_2 \\ f_3 \end{bmatrix}_{(t)} \tag{A.1}$$

with the transition matrix:

$$M = \begin{bmatrix} 1-k & \frac{1-k}{2} & 0 \\ 0 & \frac{1-k}{2} & w_{\mathrm{X^*Y}} \frac{1-k^*}{2} \\ 0 & \frac{k}{2} & w_{\mathrm{X^*Y}} \frac{k^*}{2} \end{bmatrix} \tag{A.2}$$

$m_{ij}$ elements describe the number of females of genotype $i$ in the progeny of a female of genotype $j$.

The study of the eigensystem of the transition matrix provides the conditions for stability: the eigenvector associated to the greatest eigenvalue gives the relative numbers of the three types of females at equilibrium ($\hat{f}_1$, $\hat{f}_2$, $\hat{f}_3$), and the stability conditions can be obtained by comparing non-zero eigenvalues as follows:

The two largest eigenvalues are:
$\lambda_1 = 1 - k$ and $\lambda_2 = \dfrac{1}{4}(1 - k + k^* w_{\mathrm{X^*Y}} + \Delta)$, where $\Delta = \sqrt{4(k - k^*)w_{\mathrm{X^*Y}} + (1 - k + k^* w_{\mathrm{X^*Y}})^2}$
with associated right eigenvectors:
$v_1 = (1, 0, 0)$ and $v_2 = \left( \dfrac{(k-1)(k + k^* w_{\mathrm{X^*Y}} + \Delta - 1)}{k(3k + k^* w_{\mathrm{X^*Y}} - \Delta - 3)}, -\dfrac{k + k^* w_{\mathrm{X^*Y}} + \Delta - 1}{2k}, 1 \right)$

If $\lambda_1$ is the leading eigenvalue, all females are XX at equilibrium ($\hat{f}_1 = 1$), the X* is therefore absent. If $\lambda_2$ is the leading eigenvalue, the three types of females are present at equilibrium, (their relative number is given by the elements of $v_2$). Therefore, the polygenic sex determination system is stable for $\lambda_1 < \lambda_2$, *i.e.*:

$$w_{\mathrm{X^*Y}} > \frac{(1-k)^2}{\frac{k}{2} - k^*(k - \frac{1}{2})} \tag{A.3}$$

The polygenic sex determination system is stable if the fertility of X*Y females ($w_{\mathrm{X^*Y}}$) exceeds a threshold value that depends on both $k$ and $k^*$ (the transmission ratio of male sex chromosomes crossed with XX and XX* females on the one hand and X*Y females on the other). This threshold value decreases with increasing values of $k$; and with $k > 0.5$, it is always smaller than one (see fig. 1 in main text); in other words, the more males transmit their Y chromosome in crosses with XX and XX* females, the more likely is the X* maintained at equilibrium (it is maintained for a wider range of $w_{\mathrm{X^*Y}}$ values), and with Y chromosome drive, the X* can be maintained despite a lower fertility of X*Y females ($w_{\mathrm{X^*Y}} < 1$). The impact of $k^*$ on stability depends on whether $w_{\mathrm{X^*Y}}$ is smaller or greater than one. With $w_{\mathrm{X^*Y}} < 1$, stability is made easier by smaller values of $k^*$, as it allows to reduce the production of the less fit X*Y females. With $w_{\mathrm{X^*Y}} > 1$, stability is made easier by larger values of $k^*$, as it allows the production of more X*Y females.

#### Equilibrium frequencies of males and females

The recursions (A.1) can also be written in equation form, including males:

$$\begin{aligned}
f_1' &= f_1(1-k) + f_2\frac{1-k}{2} \\
f_2' &= f_2\frac{1-k}{2} + f_3 w_{X^*Y}\frac{1-k^*}{2} \\
f_3' &= f_2\frac{k}{2} + f_3 w_{X^*Y}\frac{k^*}{2} \\
m_1' &= f_1 k + f_2\frac{k}{2} + f_3 w_{X^*Y}\frac{1-k^*}{2}
\end{aligned} \tag{A.4}$$

where $f_i$' and $m_j$' are the numbers of females of genotype $i$ and males of genotype $j$ at the next generation.

The frequencies of the $i$-th female genotype and $j$-th male genotype are written $x_i$ and $y_j$ (with $\sum_i x_i + \sum_j y_j = 1$), XX, XX* and X*Y females being in frequencies ($x_1$, $x_2$, $x_3$) and XY males in frequency ($y_1$), and are given by:

$$x_i = \frac{f_i}{\sum_i f_i + \sum_j m_j} \text{ and } y_j = \frac{m_j}{\sum_i f_i + \sum_j m_j}$$

Equilibrium frequencies ($\hat{y}_1$, $\hat{x}_1$, $\hat{x}_2$, $\hat{x}_3$) can be inferred by implementing the relative number of females at equilibrium ($\hat{f}_1$, $\hat{f}_2$ and $\hat{f}_3$) in these equations (see fig. S1).

#### Dual advantage of the X*

A Y-chromosome drive ($k > 0.5$) provides an advantage to the X* over the X by facilitating its maintenance even if females suffer from lower fertility (eq. (A.3)). We have two non-mutually-exclusive hypotheses to explain this advantage: (i) the X* is favoured by sex ratio selection, as it allows producing more females, the rarer sex in the presence of a Y chromosome drive, and (ii) the X* is advantageous because it resists Y-drive: X*Y females transmit their sex chromosomes equally, as opposed to males. To test these hypotheses, we slightly modified the model. For simplicity, we set $k = k^*$, and modified the transmission ratio of sex chromosomes in X*Y females: instead of having a Mendelian transmission, they transmit their Y at a rate $k$, like male. This mimics a situation in which the X* does not resist Y-drive.

The dynamics of the model is given by the recursions (A.1), this time with transition matrix:

$$M = \begin{bmatrix} 1-k & \frac{1-k}{2} & 0 \\ 0 & \frac{1-k}{2} & w_{X^*Y}(1-k)^2 \\ 0 & \frac{k}{2} & w_{X^*Y}k(1-k) \end{bmatrix} \tag{A.5}$$

The two largest eigenvalues are: $\lambda_1 = 1-k$ and $\lambda_2 = \frac{1-k}{2}(1 + 2kw_{X^*Y})$
with associated eigenvectors: $v_1 = (1,0,0)$ and $v_2 = \left(\frac{1-k}{2kw_{X^*Y}-k}, \frac{(1-k)(2kw_{X^*Y}-1)}{2kw_{X^*Y}-k}, \frac{k(2kw_{X^*Y}-1)}{2kw_{X^*Y}-k}\right)$

Therefore, the system is stable if $\lambda_1 < \lambda_2$ *i.e.*:

$$w_{X^*Y} > \frac{1}{2k} \tag{A.6}$$

$\frac{1}{2k}$ is inversely proportional to $k$, and for $k > 0.5$, it is always smaller than 1. This means that Y-drive would promote the spread of an X* even if it did not resist drive.

However, comparing (A.6) to (A.3) with $k = k^*$: $w_{X^*Y} > \frac{1-k}{k}$, shows that the X* can be maintained for lower $w_{X^*Y}$ values if X*Y females resist Y-drive ($\frac{1}{2k} - \frac{1-k}{k} = 1 - \frac{1}{2k}$, always positive for $k > 0.5$). This shows that the advantage of the X* has two components: it is advantaged even if it does not resist drive (likely thanks to sex ratio selection), and gets an even greater advantage if it does resist drive.

### Appendix B - Paths to the evolution of a polygenic sex determination system with conditional sex chromosome drive

The polygenic sex determination system with conditional drive of male sex chromosome found in the African Pygmy mouse evolved from a standard male heterogametic system with no sex chromosome drive. Here we present the analyses focusing on evaluating the different evolutionary scenarios (presented in the main text, fig. 2) proposed to explain this transition. These scenarios are divided into several steps, each step corresponding to the spread a single mutation. The analyses consist in defining the conditions allowing for the spread (and fixation when relevant) of the mutation at each step. For clarity, we grouped analyses based on the type of chromosome that carries the mutation involved (X*, Y, autosome), and we use the index of each step (a1, a2...) for easy reference.

#### Invasion of a mutant X*

All models focusing on the conditions allowing the invasion of a rare X* chromosome are linear, because the X* is only ever carried by females. The conditions allowing a rare X* to invade can therefore be inferred from a standard stability analysis (see Appendix A for details on the procedure).

##### Steps b1, a2 and a2'

Conditions for invasion of a rare feminizing X* chromosome at steps b1, a2 and a2' can be derived from the equation (A.3), which provides the stability conditions of the X* in a system with conditional drive of male sex chromosomes ($k \neq k^*$):

- **Step b1: invasion of an X* in a XX/XY system with no sex chromosome drive ($k = k^* = \frac{1}{2}$).** Replacing $k$ and $k^*$ by $\frac{1}{2}$ in (A.3) provides the condition for invasion:

$$w_{\mathrm{X^*Y}} > 1 \tag{B.1}$$

- **Step a2: invasion of an X* in a XX/XY system with a pre-existing non-conditional sex chromosome drive ($k^* = k$).** Replacing $k^*$ by $k$ in equation (A.3) provides the condition for invasion:

$$w_{\mathrm{X^*Y}} > \frac{1-k}{k} \tag{B.2}$$

- **Step a2': invasion of an X* which modifies the transmission of male sex chromosomes in crosses with X*Y females, in a system with a pre-existing sex chromosome drive ($k^* \neq k$).** The condition for invasion is directly given by (A.3) :

$$w_{\mathrm{X^*Y}} > \frac{(1-k)^2}{\frac{k}{2} - k^*(k - \frac{1}{2})} \tag{B.3}$$

##### Step 3 - invasion of a X* that modifies drive in XY x X*Y crosses, in a XX,XX*,X*Y/XY system with unconditional male sex chromosome drive

In this case, the system is also linear but a little more complex: the mutant X* that affects male sex chromosome drive in crosses with X*Y females, thereafter called $\mathrm{X^{*d}}$, appears in a system in which there is (i) an unconditional drive of male sex chromosome and (ii) a resident X* (that has no effect on sex chromosome transmission, as opposed to $\mathrm{X^{*d}}$).

The model is composed of one type of males: XY, and five types of females: XX, XX*, $\mathrm{XX^{*d}}$, X*Y and $\mathrm{X^{*d}Y}$, in numbers ($f_1$ ... $f_5$). The transmission ratio of male sex chromosomes depends on female genotype: $k$ in crosses with XX, XX*, $\mathrm{XX^{*d}}$ and X*Y females, and $k^*$ in crosses with $\mathrm{X^{*d}Y}$ females. Females have different fertilities ($w_{\mathrm{XX}} = w_{\mathrm{XX^*}} = w_{\mathrm{XX^{*d}}} = 1$ and $w_{\mathrm{X^*Y}} = w_{\mathrm{X^{*d}Y}}$), and we assume that the driving allele is not costly.

The dynamics of the model is given by the recursions:

$$\begin{bmatrix} f_1 \\ \vdots \\ f_5 \end{bmatrix}_{(t+1)} = M \begin{bmatrix} f_1 \\ \vdots \\ f_5 \end{bmatrix}_{(t)} \tag{B.4}$$

With the transition matrix:

$$T_3 = \begin{bmatrix} 1-k & \frac{1-k}{2} & \frac{1-k}{2} & 0 & 0 \\ 0 & \frac{1-k}{2} & 0 & w_{\mathrm{X^*Y}}\frac{1-k}{2} & 0 \\ 0 & 0 & \frac{1-k}{2} & 0 & w_{\mathrm{X^*Y}}\frac{1-k^*}{2} \\ 0 & \frac{k}{2} & 0 & w_{\mathrm{X^*Y}}\frac{k}{2} & 0 \\ 0 & 0 & \frac{k}{2} & 0 & w_{\mathrm{X^*Y}}\frac{k^*}{2} \end{bmatrix} \tag{B.5}$$

The three leading eigenvalues are: $\lambda_1 = 1-k$, $\lambda_2 = \frac{1}{2}(1+k(w_{\mathrm{X^*Y}}-1))$ and $\lambda_3 = \frac{1}{4}(1-k+k^*w_{\mathrm{X^*Y}} + \sqrt{k^2-2k(1+(k^*-2)w_{\mathrm{X^*Y}})+(k^*-1)^2})$.

According to the eigenvectors (not shown here), the $\mathrm{X^{*d}}$ can only invade if $\lambda_3$ is the leading eigenvalue (*i.e.*, if it is greater than $\lambda_1$ and $\lambda_2$).

- $\lambda_3 > \lambda_2$ if $w_{\mathrm{X^*Y}} > \frac{(1-k)^2}{\frac{k}{2}-k^*(k-\frac{1}{2})}$ (condition for stability of the X* in a polygenic system with conditional drive of male sex chromosomes (eq. (A.3) in Appendix A)).
- $\lambda_3 > \lambda_1$ if (i) $k^* > k$ (when $w_{\mathrm{X^*Y}} > 1$); (ii) $k^* < k$ (when $\frac{1-k}{k} < w_{\mathrm{X^*Y}} < 1$).

The condition for invasion of a driving X* depends on the fertility of X*Y females: if it is superior to that of XX and XX* females ($w_{\mathrm{X^*Y}} > 1$), the original X* will be replaced provided that the new one increases the strength of Y drive ($k^* > k$), as it gains a fitness advantage from producing more of the fitter X*Y females. Alternatively, if X*Y females have a lower fertility ($w_{\mathrm{X^*Y}} < 1$), then the $\mathrm{X^{*d}}$ will invade provided it decreases Y drive ($k^* < k$), as it reduces this reduces the production of X*Y females.

### Invasion of a Y-linked mutant

Models focusing on the the spread of mutant Y-linked alleles affecting the transmission ratio of male sex chromosomes are non-linear, because the mutant allele is carried by both males and females (with the exception of step a1). The conditions for invasion of such mutants are defined from stability analyses for non-linear models with multiple variables [S1]. The method relies on the calculation of the eigenvalues of the Jacobian matrix associated with the system of equations for the equilibrium of interest (mutant allele absent). If the leading eigenvalue is greater than one: the equilibrium is unstable *i.e.*, the mutant allele will increase in frequency when rare. Because a successful invasion does not entail that the mutant allele goes to fixation, we also analyse the Jacobian matrix of the equilibrium point where the mutant allele is fixed: if all its eigenvalues are smaller than one: the system is stable against the spread of wild-type alleles *i.e.*, assuming the mutant allele is able to spread and reaches high frequency, it goes to fixation.

#### Step a1 - invasion of a driving Y in a XX/XY system with no sex chromosome drive

The dynamics of the spread of a Y chromosome driver in a simple male heterogametic sex determination system was explored analytically by Hamilton [S2], who showed that alleles inherited in a biased manner can spread in a population despite not providing a fitness advantage to its bearers. In other words, a driving Y chromosome will spread and go to fixation as long as it favours its own transmission ($k > 0.5$).

#### Steps b2, b2' and 3

These three steps involve the invasion of a Y-linked driver of male sex chromosomes in a polygenic sex determination system. In step b2, a non-conditional driver spreads in a system with no pre-existing sex chromosome drive. In step b2', a conditional driver spreads in a system with no pre-existing drive. In step 3, the Y-linked driver modifies a pre-existing, non-conditional drive in crosses with X*Y females. The conditions for invasion and fixation of the mutant allele in these three steps can be derived by analysing simplified versions of the following general model:

- The model

The alleles considered are : X, X*, Y, $Y^d$ (the driving allele). There are two types of males: XY and $XY^d$ and four types of females: XX, XX*, X*Y and $X^*Y^d$, in numbers $m_j$ ($m_1$, $m_2$) and $f_i$ ($f_1 ... f_4$) respectively. We assume that the driving allele is not costly: XY and $XY^d$ have the same fertility, and all sex reversed females (X*Y and $X^*Y^d$) have a fertility $w_{X^*Y}$. In line with empirical observations, the transmission ratio of female sex chromosomes is mendelian. In males: the Y allele has a transmission ratio of $k_a$ in crosses with XX, XX* and X*Y females, and $k_b$ in crosses with $X^*Y^d$ females; the $Y^d$ allele has a transmission ratio of $k_c$ in crosses with XX, XX* and X*Y females, and $k_d$ in crosses with $X^*Y^d$ females (table B1).

| crosses | | offspring genotypes | | | | | |
|---|---|---|---|---|---|---|---|
| male | female | XY | $XY^d$ | XX | XX* | X*Y | $X^*Y^d$ |
| XY | XX | $k_a$ | 0 | $1-k_a$ | 0 | 0 | 0 |
| XY | XX* | $\frac{k_a}{2}$ | 0 | $\frac{1-k_a}{2}$ | $\frac{1-k_a}{2}$ | $\frac{k_a}{2}$ | 0 |
| XY | X*Y | $\frac{1-k_a}{2}$ | 0 | 0 | $\frac{1-k_a}{2}$ | $\frac{k_a}{2}$ | 0 |
| XY | $X^*Y^d$ | 0 | $\frac{1-k_b}{2}$ | 0 | $\frac{1-k_b}{2}$ | $\frac{k_b}{2}$ | 0 |
| $XY^d$ | XX | 0 | $k_c$ | $1-k_c$ | 0 | 0 | 0 |
| $XY^d$ | XX* | 0 | $\frac{k_c}{2}$ | $\frac{1-k_c}{2}$ | $\frac{1-k_c}{2}$ | 0 | $\frac{k_c}{2}$ |
| $XY^d$ | X*Y | $\frac{1-k_c}{2}$ | 0 | 0 | $\frac{1-k_c}{2}$ | 0 | $\frac{k_c}{2}$ |
| $XY^d$ | $X^*Y^d$ | 0 | $\frac{1-k_d}{2}$ | 0 | $\frac{1-k_d}{2}$ | 0 | $\frac{k_d}{2}$ |

**Table B1.** Offspring produced in all possible crosses in a XX,XX*,X*Y/XY system with a mutant $Y^d$ that modifies the transmission ratio of male sex chromosomes.

The dynamics of the model can be described by the following recursions:

$$
\begin{aligned}
f_1' &= \frac{1}{\sum_j m_j}\left[(m_1(1-k_a)+m_2(1-k_c))\left(f_1+\frac{f_2}{2}\right)\right]\\
f_2' &= \frac{1}{\sum_j m_j}\left[m_1\left(\frac{1-k_a}{2}(f_2+f_3w_{X^*Y})+\frac{1-k_b}{2}f_4w_{X^*Y}\right)+m_2\left(\frac{1-k_c}{2}(f_2+f_3w_{X^*Y})+\frac{1-k_d}{2}f_4w_{X^*Y}\right)\right]\\
f_3' &= \frac{1}{\sum_j m_j}\left[m_1\left(\frac{k_a}{2}(f_2+f_3w_{X^*Y})+\frac{k_b}{2}f_4w_{X^*Y}\right)\right]\\
f_4' &= \frac{1}{\sum_j m_j}\left[m_2\left(\frac{k_c}{2}(f_2+f_3w_{X^*Y})+\frac{k_d}{2}f_4w_{X^*Y}\right)\right]\\
m_1' &= \frac{1}{\sum_j m_j}\left[m_1\left(k_a(f_1+\frac{f_2}{2})+\frac{1-k_a}{2}f_3w_{X^*Y}\right)+m_2\frac{1-k_c}{2}f_3w_{X^*Y}\right]\\
m_2' &= \frac{1}{\sum_j m_j}\left[m_1\frac{1-k_b}{2}f_4w_{X^*Y}+m_2\left(k_c\left(f_1+\frac{f_2}{2}\right)+\frac{1-k_d}{2}f_4w_{X^*Y}\right)\right]
\end{aligned}
\tag{B.6}
$$

where $f_i'$ and $m_j'$ are the number of females of genotype $i$ and males of genotype $j$ at the next generation.

The intra-sex frequencies of the $i$-th female genotype and $j$-th male genotype are written $x'_i$ and $y'_j$ (with $\sum_i x'_i = 1$, $\sum_j y'_j = 1$) and are given by: $x'_i = \dfrac{f'_i}{\sum_i f'_i}$ and $y'_j = \dfrac{m'_j}{\sum_j m'_j}$.

The conditions for invasion or fixation of a driving $Y^d$ allele can be obtained by studying the eigenvalues of the Jacobian matrix (B.7) associated with the system of equations (B.6) for the equilibrium of interest.

$$J = \begin{bmatrix} \frac{\partial x'_1}{\partial \hat{x}_1} & \frac{\partial x'_1}{\partial \hat{x}_2} & \frac{\partial x'_1}{\partial \hat{x}_3} & \frac{\partial x'_1}{\partial \hat{x}_4} & \frac{\partial x'_1}{\partial \hat{y}_1} & \frac{\partial x'_1}{\partial \hat{y}_2} \\ \frac{\partial x'_2}{\partial \hat{x}_1} & \frac{\partial x'_2}{\partial \hat{x}_2} & \frac{\partial x'_1}{\partial \hat{x}_3} & \frac{\partial x'_2}{\partial \hat{x}_4} & \frac{\partial x'_2}{\partial \hat{y}_1} & \frac{\partial x'_2}{\partial \hat{y}_2} \\ \frac{\partial x'_3}{\partial \hat{x}_1} & \frac{\partial x'_3}{\partial \hat{x}_2} & \frac{\partial x'_3}{\partial \hat{x}_3} & \frac{\partial x'_3}{\partial \hat{x}_4} & \frac{\partial x'_3}{\partial \hat{y}_1} & \frac{\partial x'_3}{\partial \hat{y}_2} \\ \frac{\partial x'_4}{\partial \hat{x}_1} & \frac{\partial x'_4}{\partial \hat{x}_2} & \frac{\partial x'_4}{\partial \hat{x}_3} & \frac{\partial x'_4}{\partial \hat{x}_4} & \frac{\partial x'_4}{\partial \hat{y}_1} & \frac{\partial x'_4}{\partial \hat{y}_2} \\ \frac{\partial y'_1}{\partial \hat{x}_1} & \frac{\partial y'_1}{\partial \hat{x}_2} & \frac{\partial y'_1}{\partial \hat{x}_3} & \frac{\partial y'_1}{\partial \hat{x}_4} & \frac{\partial y'_1}{\partial \hat{y}_1} & \frac{\partial y'_1}{\partial \hat{y}_2} \\ \frac{\partial y'_2}{\partial \hat{x}_1} & \frac{\partial y'_2}{\partial \hat{x}_2} & \frac{\partial y'_2}{\partial \hat{x}_3} & \frac{\partial y'_2}{\partial \hat{x}_4} & \frac{\partial y'_2}{\partial \hat{y}_1} & \frac{\partial y'_2}{\partial \hat{y}_2} \end{bmatrix} \tag{B.7}$$

- Invasion of the mutant allele

The equilibrium of interest is the one where $Y^d$ is absent: at this equilibrium, all males are XY ($\hat{y}_1 = 1$, $\hat{y}_2 = 0$), $X^*Y^d$ females are absent ($\hat{x}_4 = 0$), and the frequencies of XX, XX* and X*Y females ($\hat{x}_1$, $\hat{x}_2$, $\hat{x}_3$) are given by the eigenvectors associated to the leading eigenvalue of transition matrix (A.2) with $k = k^* = k_a$. The leading eigenvalue of this matrix is: $\lambda = \dfrac{w_{X^*Y}(k_a + 1) - 1}{2}$, and female equilibrium frequencies are worth:

$$\hat{x}_1 = \frac{(k_a - 1)^2}{k_a(w_{X^*Y} + k_a - 1)},\ \hat{x}_2 = \frac{(k_a(w_{X^*Y} - 1) - 1)(1 - k_a)}{k_a(w_{X^*Y} + k_a - 1)} \text{ and } \hat{x}_3 = \frac{k_a(w_{X^*Y} + k_a) - 1}{k_a + w_{X^*Y} - 1}$$

These values, along with the intra-sex frequencies of females ($x'_i$) and males ($y'_j$), are implemented in the Jacobian (B.7). The resulting matrix, thereafter referred to as $J_{invY}$, is shown on the next page.

- Fixation of the mutant allele

The equilibrium of interest is the one where $Y^d$ is fixed: at this equilibrium, all males are $XY^d$ ($\hat{y}_1 = 0$, $\hat{y}_2 = 1$), X*Y females are absent ($\hat{x}_3 = 0$), and the frequencies of XX, XX* and $X^*Y^d$ females ($\hat{x}_1$, $\hat{x}_2$, $\hat{x}_4$) are given by the eigenvector associated to the leading eigenvalue of transition matrix (A.2) with $k = k_c$ and $k* = k_d$. The leading eigenvalue of this matrix is: $\lambda_2 = \dfrac{1}{4}(1 - k_c + k_d w_{X^*Y} + \Delta)$, where $\Delta = \sqrt{4(k_c - k_d)w_{X^*Y} + (1 - k_c + k_d w_{X^*Y})^2}$

Female equilibrium frequencies are worth:

$$\hat{x}_1 = \frac{(k_c - 1)(k_c - 1 + k_d w_{X^*Y} - \Delta)}{k_c(3(k_c - 1) + w_{X^*Y}(2 - k_d) + \Delta)}$$

$$\hat{x}_2 = \frac{-(k_c - 1 + k_d w_{X^*Y} - \Delta)(3(k_c - 1) + k_d w_{X^*Y} + \Delta)}{2k_c(3(k_c - 1) + w_{X^*Y}(2 - k_d) + \Delta)}$$

$$\hat{x}_3 = \frac{3(k_c - 1) + k_d w_{X^*Y} + \Delta}{3(k_c - 1) + w_{X^*Y}(2 - k_d) + \Delta}.$$

These values, along with the intra-sex frequencies of females ($x'_i$) and males ($y'_j$), are implemented in the Jacobian (B.7). The resulting matrix, thereafter referred to as $J_{fixY}$, is too large and complex to show here.

$$
J_{\mathrm{invY}} = \begin{bmatrix}
\frac{2(1-k_a)(k_a w_{\mathrm{X*Y}}+k_a-1)}{k_a(1+k_a(w_{\mathrm{X*Y}}-1))(k_a+w_{\mathrm{X*Y}}-1)} & \frac{(1-k_a)(k_a(2+w_{\mathrm{X*Y}})-2)}{k_a(1+k_a(w_{\mathrm{X*Y}}-1))(k_a+w_{\mathrm{X*Y}}-1)} & \frac{(1-k_a)^2 w_{\mathrm{X*Y}}}{k_a(1+k_a(w_{\mathrm{X*Y}}-1))(k_a+w_{\mathrm{X*Y}}-1)} & \frac{(1-k_a)^2 w_{\mathrm{X*Y}}}{k_a(1+k_a(w_{\mathrm{X*Y}}-1))(k_a+w_{\mathrm{X*Y}}-1)} & 0 & \frac{(1-k_a)(k_a-k_c)(k_a w_{\mathrm{X*Y}}+k_a-1)}{k_a^2(k_a+w_{\mathrm{X*Y}}-1)^2} \\
\frac{2(1-k_a)(k_a w_{\mathrm{X*Y}}+k_a-1)}{k_a(1+k_a(w_{\mathrm{X*Y}}-1))(k_a+w_{\mathrm{X*Y}}-1)} & \frac{(1-k_a)^2(k_a(2+w_{\mathrm{X*Y}})-2)}{k_a(1+k_a(w_{\mathrm{X*Y}}-1))(k_a+w_{\mathrm{X*Y}}-1)} & \frac{(1-k_a)^3 w_{\mathrm{X*Y}}}{k_a(1+k_a(w_{\mathrm{X*Y}}-1))(k_a+w_{\mathrm{X*Y}}-1)} & \frac{w_{\mathrm{X*Y}}(1+k_a^2(2-k_b+w_{\mathrm{X*Y}})+k_a(-3+k_b-k_b w_{\mathrm{X*Y}}))}{k_a(1+k_a(w_{\mathrm{X*Y}}-1))(k_a+w_{\mathrm{X*Y}}-1)} & 0 & \frac{(k_a-k_c)(k_a w_{\mathrm{X*Y}}+k_a-1)^2}{k_a^2(k_a+w_{\mathrm{X*Y}}-1)^2} \\
\frac{2(k_a-1)(k_a w_{\mathrm{X*Y}}+k_a-1)}{(1+k_a(w_{\mathrm{X*Y}}-1))(k_a+w_{\mathrm{X*Y}}-1)} & \frac{(k_a-1)(k_a(2+w_{\mathrm{X*Y}})-2)}{(1+k_a(w_{\mathrm{X*Y}}-1))(k_a+w_{\mathrm{X*Y}}-1)} & \frac{(1-k_a)^2 w_{\mathrm{X*Y}}}{(1+k_a(w_{\mathrm{X*Y}}-1))(k_a+w_{\mathrm{X*Y}}-1)} & \frac{w_{\mathrm{X*Y}}(k_b(w_{\mathrm{X*Y}}-1)-k_a(1-k_b+w_{\mathrm{X*Y}})+1}{(1+k_a(w_{\mathrm{X*Y}}-1))(k_a+w_{\mathrm{X*Y}}-1)} & 0 & \frac{(1-k_a(w_{\mathrm{X*Y}}+1))((k_a-1)k_c+k_a w_{\mathrm{X*Y}})}{k_a(k_a+w_{\mathrm{X*Y}}-1)^2} \\
0 & 0 & 0 & 0 & 0 & \frac{k_c(k_a(w_{\mathrm{X*Y}}+1)-1)}{k_a(k_a+w_{\mathrm{X*Y}}-1)} \\
0 & 0 & 0 & \frac{(1-k_b)w_{\mathrm{X*Y}}(k_a+w_{\mathrm{X*Y}}-1)}{(k_a-1)(1-w_{\mathrm{X*Y}}+k_a(-1+2w_{\mathrm{X*Y}}+w_{\mathrm{X*Y}}{}^2))} & 0 & \frac{k_c(k_a(w_{\mathrm{X*Y}}-1)-1)}{k_a(1-w_{\mathrm{X*Y}}+k_a(-1+2w_{\mathrm{X*Y}}+w_{\mathrm{X*Y}}{}^2))} \\
0 & 0 & 0 & \frac{(k_b-1)w_{\mathrm{X*Y}}(k_a+w_{\mathrm{X*Y}}-1)}{(k_a-1)(1-w_{\mathrm{X*Y}}+k_a(-1+2w_{\mathrm{X*Y}}+w_{\mathrm{X*Y}}{}^2))} & 0 & \frac{k_c(k_a(w_{\mathrm{X*Y}}+1)-1)}{k_a(1-w_{\mathrm{X*Y}}+k_a(-1+2w_{\mathrm{X*Y}}+w_{\mathrm{X*Y}}{}^2))}
\end{bmatrix}
\tag{B.8}
$$

The complexity of the Jacobian matrices $J_{invY}$ and $J_{fixY}$ precludes the analysis of their eigenvalues, and therefore to derive general analytical conditions for invasion and fixation of Y-linked driving alleles. As a workaround, we analyse simplified versions of the Jacobians, that conform to each of the three steps 2b, 2b' and 3. These simplified Jacobians were obtained by replacing $k_a$, $k_b$, $k_c$ and $k_d$ by $\frac{1}{2}$, $k$ or $k^*$ as shown in table B2, which also displays a summary of invasion and fixation conditions. The full analyse of the eigenvalues of $J_{invY}$ and $J_{fixY}$ for each case is detailed below.

| | **step b2** | **step b2'** | **step 3** |
|---|---|---|---|
| | $k_a = k_b = \frac{1}{2}$ | $k_a = k_b = \frac{1}{2}$ | $k_a = k_c = k$ |
| | $k_c = k_d = k$ | $k_c = k$ and $k_d = k^*$ | $k_b = k_d = k^*$ |
| **X* stability** | $w_{\mathrm{X^*Y}} > 1$ | $w_{\mathrm{X^*Y}} > 1$ | $w_{\mathrm{X^*Y}} > \frac{1-k}{k}$ |
| **Condition for invasion** | $k > \frac{1}{2}$ | $k > \frac{1}{2}$ | $k^* < k$ |
| **Condition for fixation** | $k > \frac{1}{2}$ | $k^* < \frac{k^3(2-2w_{\mathrm{X^*Y}})+k^2(3w_{\mathrm{X^*Y}}{}^2+6w-5)+k(4-5w_{\mathrm{X^*Y}}-w_{\mathrm{X^*Y}})+w_{\mathrm{X^*Y}}-1}{2kw_{\mathrm{X^*Y}}(kw_{\mathrm{X^*Y}}+k-1)}$ † | $k^* < k$ |

**Table B2. Conditions for invasion and fixation of mutant Y-linked male sex chromosome drivers.** For each step (b2, b2' and 3), after defining the condition for X* stability in the absence of the mutant allele, conditions for invasion and fixation were defined by analysing the eigenvalues of simplified versions of $J_{invY}$ and $J_{fixY}$, in which $k_a$, $k_b$, $k_c$ and $k_d$ were replaced by the appropriate numerical value or substituted by $k$ or $k^*$. †: see fig. B1.

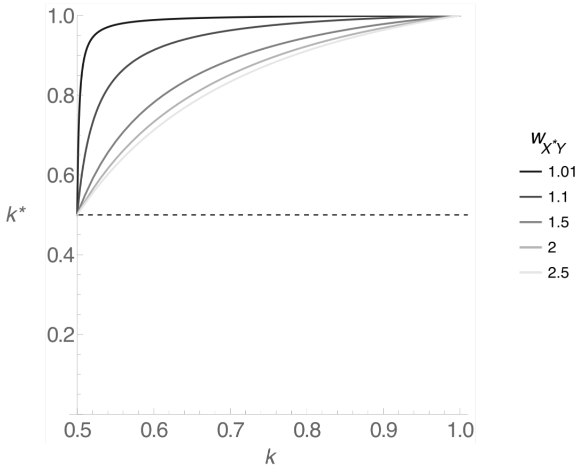

**Figure B1. Conditions for fixation of a conditionally driving $\mathrm{Y^d}$ in a XX,XX*,X*Y/XY system with no pre-existing sex chromosome drive** for different values of $w_{\mathrm{X^*Y}}$. The space below each line shows the $k$ x $k^*$ parameter space in which the leading eigenvalue of the Jacobian matrix associated to the equilibrium where $\mathrm{Y^d}$ is fixed is smaller than one, *i.e*, the parameter space in which a spreading $\mathrm{Y^d}$ goes to fixation.

- **step b2: invasion of a Y-linked driver of male sex chromosomes $Y^d$ (unconditional) in a system with no pre-existing sex chromosome drive ($k_a = k_b = \frac{1}{2}$ and $k_c = k_d = k$)**

**invasion of $Y^d$**: Assuming the condition for stability of the X* chromosome at the equilibrium point where $Y^d$ is absent is satisfied ($w_{X^*Y} > 1$, eq. (A.3) with $k = k^* = \frac{1}{2}$), the leading eigenvalue of the $J_{invY}$ with $k_a = k_b = \frac{1}{2}$ and $k_c = k_d = k$ is:

$$\lambda = \frac{\sqrt{k\left(k(w_{X^*Y}+1)^2 + 2(w_{X^*Y}-1)w_{X^*Y}\left(w_{X^*Y}{}^2+1\right)\right)} + kw_{X^*Y} + k}{w_{X^*Y}{}^2+1}$$

$\lambda > 1$ for $k > \frac{1}{2}$, meaning that a $Y^d$ will invade if it favors its own transmission, like in a standard male heterogametic system.

**fixation of $Y^d$**: Assuming the condition for stability of the X* ($w_{X^*Y} > 1$) and invasion of the $Y^d$ allele ($k > \frac{1}{2}$) are satisfied, the leading eigenvalue of the $J_{fixY}$ with $k_a = k_b = \frac{1}{2}$ and $k_c = k_d = k$ is:

$$\lambda = \frac{1 + k(w_{X^*Y} - 1) - \sqrt{1 + 2k(w_{X^*Y} - 1)(4w_{X^*Y} + 1) + k^2(1 + w_{X^*Y}(14 - w_{X^*Y}(15 + 16w_{X^*Y}))) + 8k^3 w_{X^*Y}(w_{X^*Y}{}^3 + 3w_{X^*Y}{}^2 + w_{X^*Y} - 1)}}{4k(w_{X^*Y} - 1 + k(w_{X^*Y}()2 + w_{X^*Y}) - 1)}$$

$\lambda < 1$ for $k > \frac{1}{2}$: a $Y^d$ chromosome in high frequency will fix if $k > \frac{1}{2}$.

- **step b2': invasion of a Y-linked conditional driver of male sex chromosomes $Y^d$ in a system with no pre-existing sex chromosome drive ($k_a = k_b = \frac{1}{2}$, $k_c = k$ and $k_d = k^*$)**

**invasion of $Y^d$**: Assuming the condition for stability of the X* chromosome at the equilibrium point where $Y^d$ is absent is satisfied ($w_{X^*Y} > 1$, eq. (A.3) with $k = k^* = \frac{1}{2}$), the leading eigenvalue of $J_{invY}$, with $k_a = k_b = \frac{1}{2}$, $k_c = k$, $k_d = k^*$ is:

$$\lambda = \frac{k + kw_{X^*Y} - i\sqrt{k}\sqrt{(1-w_{X^*Y})^2 - 2(w_{X^*Y}-1)w_{X^*Y}(w_{X^*Y}{}^2+1) - k}}{w_{X^*Y}{}^2+1}$$

$\lambda > 1$ for $k > \frac{1}{2}$: a driving $Y^d$ will invade if it favors its own transmission, like in a standard male heterogametic system and step b2.

**fixation of $Y^d$**: Assuming the condition for stability of the X* ($w_{X^*Y} > 1$) and invasion of the $Y^d$ allele ($k > \frac{1}{2}$) are satisfied, the leading eigenvalue of the $J_{fixY}$ with $k_a = k_b = \frac{1}{2}$, $k_c = k$, $k_d = k^*$ is:

$$\lambda = \frac{\sqrt{k-1}\sqrt{8k^3(k^*-1)w_{X^*Y}{}^4 + 8(k-1)k^2 w_{X^*Y}{}^3(k+2k^*-2) + (k-1)kw_{X^*Y}{}^2(8(k-1)k^*-7k+8) - 2(k-1)^2k(4k-3)w_{X^*Y} + (k-1)^3} + k(k(w_{X^*Y}-1) - w_{X^*Y}+2) - 1}{4k(k(k^*-1)w_{X^*Y}{}^2 + (k-1)w_{X^*Y}(k+k^*-1) - (k-1)^2}$$

$\lambda < 1$ for $k^* < \frac{k^3(2-2w_{X^*Y}) + k^2(3w_{X^*Y}{}^2+6w-5) + k(4-5w_{X^*Y}-w_{X^*Y}) + w_{X^*Y}-1}{2kw_{X^*Y}(kw_{X^*Y}+k-1)}$: fixation of the conditionally driving $Y^d$ depends on the values of $k$, $k^*$ and $w_{X^*Y}$ (fig. B1). If $k^*$ is smaller than $\frac{1}{2}$ (X-drive), a $Y^d$ chromosome in high frequency will go to fixation regardless of the value of $k$ and $w_{X^*Y}$. If $k^*$ is greater than $\frac{1}{2}$, values of $k$ close to one (strong Y-drive) and low values of $w_{X^*Y}$ will tend to prevent its fixation, resulting in the establishment of a polymorphic equilibrium with both Y and $Y^d$ segregating.

- **step 3: invasion of a Y-linked modifier of drive $Y^d$, specific to crosses with $X^*Y^d$ females in a system with a pre-existing male sex chromosome drive ($k_a = k_c = k$ and $k_b = k_d = k^*$)**

The mutant $Y^d$ acts in $X^*Y^d$ females: it causes males to transmit their Y chromosome at a rate $k^*$. It overrides the effect of the pre-existing male sex chromosome drive causing them to transmit their Y at a rate $k$ in all other crosses.

**invasion of $Y^d$**: Assuming the condition for stability of the X* chromosome at the equilibrium point where $Y^d$ is absent is satisfied ($w_{X^*Y} > \frac{1-k}{k}$, eq. (A.3) with $k = k^*$), the leading eigenvalue of $J_{invY}$ with $k_a = k_c = k$ and $k_b = k_d = k^*$ is:

$$\lambda = \frac{\sqrt{k-1}\sqrt{k(4k^2(k^*-1)w_{X^*Y}{}^4+4k(3k-2)(k^*-1)w_{X^*Y}{}^3+(k-1)w_{X^*Y}{}^2(4(k-1)k^*+(k-2)^2)-2(k-1)^2w_{X^*Y}(k+2k^*-2)+(k-1)^3)}+(k-1)\sqrt{k}(k(w_{X^*Y}-1)+1)}{2(k-1)\sqrt{k}(k(w_{X^*Y}(w_{X^*Y}+2)-1)-w_{X^*Y}+1)}$$

$\lambda > 1$ for $k^* < k$: a driving $Y^d$ will invade if it reduces the transmission ratio of males' Y chromosome in crosses with $X^*Y^d$ females.

**fixation of $Y^d$**: with $k_a = k_c = k$ and $k_b = k_d = k^*$, $J_{fixY}$ was too complex to compute eigenvalues. By replacing $k$ and $w_{X^*Y}$ by different sets of numerical values that comply with the stability conditions of the X* chromosome (ranging respectively from 0 to 1 (0.02 increment) and $\frac{1-k}{k}$ to 4 (0.02 increment)), we were able to show that a $Y^d$ allele in high frequency will fix for $k^* < k$, same condition as the invasion condition.

### Invasion of a mutant autosome

The study of the invasion of a sex chromosome driver carried by an autosome requires the introduction in models of a second locus, which harbors a standard non-distorting $a$ allele and a dominant distorter $A$ allele. This locus is independent from the sex chromosome locus, which implies that for each sex chromosome combination, there are three possible genotypes (*e.g.* XXaa, XXAa, XXAA). Similar to models that involve the evolution of a mutant Y-linked driver, all models presented here are non-linear, because the autosomal mutation is carried by both males and females. Therefore, conditions for invasion and fixation of a driving $A$ allele, can be defined from stability analyses for non-linear models with multiple variables (see previous part for details).

#### Steps b2, b2' and 3

These three steps involve the invasion of an autosomal driver of male sex chromosomes in a polygenic sex determination system. In step b2, a non-conditional driver spreads in a system with no pre-existing sex chromosome drive. In step b2', a conditional driver spreads in a system with no pre-existing drive. In step 3, the driver modifies a pre-existing, non-conditional drive in crosses with X*Y females. The conditions for invasion and fixation of the mutant allele in these three steps can be derived by analysing simplified versions of the following general model:

- The model

At the sex determining locus, the alleles considered are: Y, X and X*. The autosomal locus has two alleles: the ancestral non-driving $a$ allele, and the dominant driver of male sex chromosomes $A$. There are three types of males: XYaa, XYAa and XYAA and nine types of females: XXaa, XXAa, XXAA, XX*aa, XX*Aa, XX*AA, X*Yaa, X*YAa, X*YAA, in numbers $m_j$ ($m_1 ... m_3$) and $f_i$ ($f_1 ... f_9$) respectively. We assume that the driving allele is not costly: bearing one or two copies of $A$ has no impact on fitness. Like in other models, the fertility of X*Y females is allowed to vary: all sex reversed females (X*Yaa, X*YAa and X*YAA) have a fertility $w_{X^*Y}$. In line with empirical observations, the transmission ratio of female sex chromosomes is mendelian. In males, transmission of sex chromosomes depends on the presence or absence of allele A, and can be modified in crosses with X*Y females if they carry that mutation. In males homozygous for the non-driving allele $a$ (XYaa), the Y chromosome has a transmission ratio of $k_a$, except in crosses with X*YAa and X*YAA females, where it becomes $k_b$. In males carrying the mutant allele (XYAa and XYAA), the Y chromosome has a transmission ratio of $k_c$, except in crosses with X*YAa and X*YAA females, where it becomes $k_d$ (table B3).

| crosses | | offspring genotypes | | | | | | | | | | | |
|---|---|---|---|---|---|---|---|---|---|---|---|---|---|
| male | female | XYaa | XYAa | XYAA | XXaa | XXAa | XXAA | XX*aa | XX*Aa | XX*AA | X*Yaa | X*YAa | X*YAA |
| XYaa | XXaa | $k_a$ | 0 | 0 | $1-k_a$ | 0 | 0 | 0 | 0 | 0 | 0 | 0 | 0 |
| XYaa | XXAa | $\frac{k_a}{2}$ | $\frac{k_a}{2}$ | 0 | $\frac{1-k_a}{2}$ | $\frac{1-k_a}{2}$ | 0 | 0 | 0 | 0 | 0 | 0 | 0 |
| XYaa | XXAA | 0 | $k_a$ | 0 | 0 | $1-k_a$ | 0 | 0 | 0 | 0 | 0 | 0 | 0 |
| XYaa | XX*aa | $\frac{k_a}{2}$ | 0 | 0 | $\frac{1-k_a}{2}$ | 0 | 0 | $\frac{1-k_a}{2}$ | 0 | 0 | $\frac{k_a}{2}$ | 0 | 0 |
| XYaa | XX*Aa | $\frac{k_a}{4}$ | $\frac{k_a}{4}$ | 0 | $\frac{1-k_a}{4}$ | $\frac{1-k_a}{4}$ | 0 | $\frac{1-k_a}{4}$ | $\frac{1-k_a}{4}$ | 0 | $\frac{k_a}{4}$ | $\frac{k_a}{4}$ | 0 |
| XYaa | XX*AA | 0 | $\frac{k_a}{2}$ | 0 | 0 | $\frac{1-k_a}{2}$ | 0 | 0 | $\frac{1-k_a}{2}$ | 0 | 0 | $\frac{k_a}{2}$ | 0 |
| XYaa | X*Yaa | $\frac{1-k_a}{2}$ | 0 | 0 | 0 | 0 | 0 | $\frac{1-k_a}{2}$ | 0 | 0 | $\frac{k_a}{2}$ | 0 | 0 |
| XYaa | X*YAa | $\frac{1-k_b}{4}$ | $\frac{1-k_b}{4}$ | 0 | 0 | 0 | 0 | $\frac{1-k_b}{4}$ | $\frac{1-k_b}{4}$ | 0 | $\frac{k_b}{4}$ | $\frac{k_b}{4}$ | 0 |
| XYaa | X*YAA | 0 | $\frac{1-k_b}{2}$ | 0 | 0 | 0 | 0 | 0 | $\frac{1-k_b}{2}$ | 0 | 0 | $\frac{k_b}{2}$ | 0 |
| XYAa | XXaa | $\frac{k_c}{2}$ | $\frac{k_c}{2}$ | 0 | $\frac{1-k_c}{2}$ | $\frac{1-k_c}{2}$ | 0 | 0 | 0 | 0 | 0 | 0 | 0 |
| XYAa | XXAa | $\frac{k_c}{4}$ | $\frac{k_c}{2}$ | $\frac{k_c}{4}$ | $\frac{1-k_c}{4}$ | $\frac{1-k_c}{2}$ | $\frac{1-k_c}{4}$ | 0 | 0 | 0 | 0 | 0 | 0 |
| XYAa | XXAA | 0 | $\frac{k_c}{2}$ | $\frac{k_c}{2}$ | 0 | $\frac{1-k_c}{2}$ | $\frac{1-k_c}{2}$ | 0 | 0 | 0 | 0 | 0 | 0 |
| XYAa | XX*aa | $\frac{k_c}{4}$ | $\frac{k_c}{4}$ | 0 | $\frac{1-k_c}{4}$ | $\frac{1-k_c}{4}$ | 0 | $\frac{1-k_c}{4}$ | $\frac{1-k_c}{4}$ | 0 | $\frac{k_c}{4}$ | $\frac{k_c}{4}$ | 0 |
| XYAa | XX*Aa | $\frac{k_c}{8}$ | $\frac{k_c}{4}$ | $\frac{k_c}{8}$ | $\frac{1-k_c}{8}$ | $\frac{1-k_c}{4}$ | $\frac{1-k_c}{8}$ | $\frac{1-k_c}{8}$ | $\frac{1-k_c}{4}$ | $\frac{1-k_c}{8}$ | $\frac{k_c}{8}$ | $\frac{k_c}{4}$ | $\frac{k_c}{8}$ |
| XYAa | XX*AA | 0 | $\frac{k_c}{4}$ | $\frac{k_c}{4}$ | 0 | $\frac{1-k_c}{4}$ | $\frac{1-k_c}{4}$ | 0 | $\frac{1-k_c}{4}$ | $\frac{1-k_c}{4}$ | 0 | $\frac{k_c}{4}$ | $\frac{k_c}{4}$ |
| XYAa | X*Yaa | $\frac{1-k_c}{4}$ | $\frac{1-k_c}{4}$ | 0 | 0 | 0 | 0 | $\frac{1-k_c}{4}$ | $\frac{1-k_c}{4}$ | 0 | $\frac{k_c}{4}$ | $\frac{k_c}{4}$ | 0 |
| XYAa | X*YAa | $\frac{1-k_d}{8}$ | $\frac{1-k_d}{4}$ | $\frac{1-k_d}{8}$ | 0 | 0 | 0 | $\frac{1-k_d}{8}$ | $\frac{1-k_d}{4}$ | $\frac{1-k_d}{8}$ | $\frac{k_d}{8}$ | $\frac{k_d}{4}$ | $\frac{k_d}{8}$ |
| XYAa | X*YAA | 0 | $\frac{1-k_d}{4}$ | $\frac{1-k_d}{4}$ | 0 | 0 | 0 | 0 | $\frac{1-k_d}{4}$ | $\frac{1-k_d}{4}$ | 0 | $\frac{k_d}{4}$ | $\frac{k_d}{4}$ |
| XYAA | XXaa | 0 | $k_c$ | 0 | 0 | $1-k_c$ | 0 | 0 | 0 | 0 | 0 | 0 | 0 |
| XYAA | XXAa | 0 | $\frac{k_c}{2}$ | $\frac{k_c}{2}$ | 0 | $\frac{1-k_c}{2}$ | $\frac{1-k_c}{2}$ | 0 | 0 | 0 | 0 | 0 | 0 |
| XYAA | XXAA | 0 | 0 | $k_c$ | 0 | 0 | $1-k_c$ | 0 | 0 | 0 | 0 | 0 | 0 |
| XYAA | XX*aa | 0 | $\frac{k_c}{2}$ | 0 | 0 | $\frac{1-k_c}{2}$ | 0 | 0 | $\frac{1-k_c}{2}$ | 0 | 0 | $\frac{k_c}{2}$ | 0 |
| XYAA | XX*Aa | 0 | $\frac{k_c}{4}$ | $\frac{k_c}{4}$ | 0 | $\frac{1-k_c}{4}$ | $\frac{1-k_c}{4}$ | 0 | $\frac{1-k_c}{4}$ | $\frac{1-k_c}{4}$ | 0 | $\frac{k_c}{4}$ | $\frac{k_c}{4}$ |
| XYAA | XX*AA | 0 | 0 | $\frac{k_c}{2}$ | 0 | 0 | $\frac{1-k_c}{2}$ | 0 | 0 | $\frac{1-k_c}{2}$ | 0 | 0 | $\frac{k_c}{2}$ |
| XYAA | X*Yaa | 0 | $\frac{1-k_c}{2}$ | 0 | 0 | 0 | 0 | 0 | $\frac{1-k_c}{2}$ | 0 | 0 | $\frac{k_c}{2}$ | 0 |
| XYAA | X*YAa | 0 | $\frac{1-k_d}{4}$ | $\frac{1-k_d}{4}$ | 0 | 0 | 0 | 0 | $\frac{1-k_d}{4}$ | $\frac{1-k_d}{4}$ | 0 | $\frac{k_d}{4}$ | $\frac{k_d}{4}$ |
| XYAA | X*YAA | 0 | 0 | $\frac{1-k_d}{2}$ | 0 | 0 | 0 | 0 | 0 | $\frac{1-k_d}{2}$ | 0 | 0 | $\frac{k_d}{2}$ |

**Table B3.** Offspring produced in all possible crosses in a XX,XX*,X*Y/XY system with an autosomal mutant A that modifies the transmission ratio of male sex chromosomes.

The dynamics of the model can be described by the following recursions:

$$
\begin{aligned}
{m_1}' &= \frac{1}{\sum_j m_j}\Bigg[\left(m_1 k_a + m_2\frac{k_c}{2}\right)\left(f_1 + \frac{f_2}{2} + \frac{f_4}{2} + \frac{f_5}{4}\right) + \left(m_1\frac{1-k_a}{2} + m_2\frac{1-k_c}{2}\right) f_7 w_{\mathrm{X^*Y}} + \\
&\qquad \left(m_1\frac{1-k_b}{4} + m_2\frac{1-k_d}{8}\right) f_8 w_{\mathrm{X^*Y}}\Bigg] \\
{m_2}' &= \frac{1}{\sum_j m_j}\Bigg[m_1\left(k_a\left(\frac{f_2}{2} + f_3 + \frac{f_5}{4} + \frac{f_6}{2}\right) + \frac{1-k_b}{2}\left(\frac{f_8}{2} + f_9\right) w_{\mathrm{X^*Y}}\right) + \\
&\qquad m_2\left(\frac{k_c}{2}\left(f_1 + f_1 + f_3 + \frac{f_3}{2} + \frac{f_5}{2} + \frac{f_6}{2}\right) + \frac{1-k_c}{4} f_7 w_{\mathrm{X^*Y}} + \frac{1-k_d}{4}(f_8 + f_9) w_{\mathrm{X^*Y}}\right) + \\
&\qquad m_3\left(k_c\left(f_1 + \frac{f_1}{2} + \frac{f_4}{2} + \frac{f_5}{4}\right) + \frac{1-k_c}{2} f_7 w_{\mathrm{X^*Y}} + \frac{1-}{4} f_8 w_{\mathrm{X^*Y}}\right)\Bigg] \\
{m_3}' &= \frac{1}{\sum_j m_j}\left[\left(\frac{m_2}{2} + m_3\right)\left(k_c\left(\frac{f_1}{2} + f_3 + \frac{f_5}{4} + \frac{f_6}{2}\right) + \frac{1-k_d}{2}\left(\frac{f_8}{2} + f_9\right) w_{\mathrm{X^*Y}}\right)\right] \\
{f_1}' &= \frac{1}{\sum_j m_j}\left[\left(m_1(1-k_a) + m_2\frac{1-k_c}{2}\right)\left(f_1 + \frac{f_1}{2} + \frac{f_4}{2} + \frac{f_5}{4}\right)\right] \\
{f_2}' &= \frac{1}{\sum_j m_j}\Bigg[m_1(1-k_a)\left(\frac{f_2}{2} + f_3 + \frac{f_5}{4} + \frac{f_6}{2}\right) + m_2\frac{1-k_c}{2}\left(f_1 + f_2 + f_3 + \frac{f_4}{2} + \frac{f_5}{2} + \frac{f_6}{2}\right) + \\
&\qquad m_3(1-k_c)\left(f_1 + \frac{f_2}{2} + \frac{f_4}{2} + \frac{f_5}{4}\right)\Bigg] \\
{f_3}' &= \frac{1}{\sum_j m_j}\left[\left(\frac{m_2}{2} + m_3\right)(1-k_c)\left(\frac{f_2}{2} + f_3 + \frac{f_5}{4} + \frac{f_6}{2}\right)\right] \\
{f_4}' &= \frac{1}{\sum_j m_j}\left[\left(m_1\frac{1-k_a}{2} + m_2\frac{1-k_c}{4}\right)\left(f_4 + \frac{f_5}{2} + f_7 w_{\mathrm{X^*Y}}\right) + \left(m_1\frac{1-k_b}{2} + m_2\frac{1-k_d}{4}\right)\frac{f_8}{2} w_{\mathrm{X^*Y}}\right] \\
{f_5}' &= \frac{1}{\sum_j m_j}\Bigg[m_1\left(\frac{1-k_a}{2}\left(\frac{f_5}{2} + f_6\right) + \frac{1-k_b}{2}\left(\frac{f_8}{2} + f_9\right) w_{\mathrm{X^*Y}}\right) + \\
&\qquad m_2\left(\frac{1-k_c}{4}(f_4 + f_5 + f_6 + f_7 w_{\mathrm{X^*Y}}) + \frac{1-k_d}{4}(f_8 + f_9) w_{\mathrm{X^*Y}}\right) + \\
&\qquad m_3\left(\frac{1-k_c}{2}\left(f_4 + \frac{f_5}{2} + f_7 w_{\mathrm{X^*Y}}\right) + \frac{1-k_d}{4} f_8 w_{\mathrm{X^*Y}}\right)\Bigg] \\
{f_6}' &= \frac{1}{\sum_j m_j}\left[\left(\frac{m_2}{2} + m_3\right)\left(\frac{1-k_c}{2}\left(\frac{f_5}{2} + f_6\right) + \frac{1-k_d}{2}\left(\frac{f_8}{2} + f_9\right) w_{\mathrm{X^*Y}}\right)\right] \\
{f_7}' &= \frac{1}{\sum_j m_j}\left[\left(m_1 k_a + m_2\frac{k_c}{2}\right)\left(\frac{f_4}{2} + \frac{f_5}{4} + \frac{f_7}{2} w_{\mathrm{X^*Y}}\right) + \left(m_1\frac{k_b}{4} + m_2\frac{k_d}{8}\right) f_8 w_{\mathrm{X^*Y}}\right] \\
{f_8}' &= \frac{1}{\sum_j m_j}\Bigg[m_1\left(\frac{k_a}{2}\left(\frac{f_5}{2} + f_6\right) + \frac{k_b}{2}\left(\frac{f_8}{2} + f_9\right) w_{\mathrm{X^*Y}}\right) + \\
&\qquad m_2\left(\frac{k_c}{4}(f_4 + f_5 + f_6 + f_7 w_{\mathrm{X^*Y}}) + \frac{k_d}{4}(f_8 + f_9) w_{\mathrm{X^*Y}}\right) + \\
&\qquad m_3\left(\frac{k_c}{2}\left(f_4 + \frac{f_5}{2} + f_7 w_{\mathrm{X^*Y}}\right) + \frac{k_d}{4} f_8 w_{\mathrm{X^*Y}}\right)\Bigg] \\
{f_9}' &= \frac{1}{\sum_j m_j}\left[\left(\frac{m_2}{2} + m_3\right)\left(k_c\left(\frac{f_5}{4} + \frac{f_6}{2}\right) + \frac{k_d}{2}\left(\frac{f_8}{2} + f_9\right) w_{\mathrm{X^*Y}}\right)\right]
\end{aligned}
\tag{B.9}
$$

where $f_{\mathrm{i}}'$ and $m_{\mathrm{j}}'$ are the number of females and males of genotype $i$ and $j$ at the next generation.

We tried obtaining the conditions for invasion and fixation of the autosomal driver of male sex chromosomes $A$ allele in the same way we did for the Y-linked driver. The Jacobian matrices associated with the system of equations (B.9) at the two equilibria of interests ($A$ absent and $a$ absent), were too complex to compute eigenvalues and derive these conditions analytically, even when simplifying the model to match the three cases b2, b2' and 3 by replacing $k_a$, $k_b$, $k_c$ and $k_d$ by $\frac{1}{2}$, $k$ or $k^*$ as shown in table B4. Instead, we performed numerical analyses of the model to obtain conditions for invasion, and ran numerical deterministic simulations to explore under which conditions the mutant alleles can go to fixation. Results of these analyses are provided in table B4, and details are given below.

| | step b2 | step b2' | step 3 |
|---|---|---|---|
| | $k_a = k_b = \frac{1}{2}$ | $k_a = k_b = \frac{1}{2}$ | $k_a = k_c = k$ |
| | $k_c = k_d = k$ | $k_c = k$ and $k_d = k^*$ | $k_b = k_d = k^*$ |
| **X* stability** | $w_{X^*Y} > 1$ | $w_{X^*Y} > 1$ | $w_{X^*Y} > \frac{1-k}{k}$ |
| **Condition for invasion** | $k > \frac{1}{2}$ | $k > \frac{1}{2}$ | $k^* < k$ |
| **Condition for fixation** | See fig. S2 | See fig. S3 | $k^* < k$ |

**Table B4. Conditions for invasion and fixation of mutant autosomal male sex chromosome drivers.** For each step (b2, b2' and 3), condition for X* stability in the absence of the mutant allele were defined based on equation (A.1).

- **step b2: invasion of an autosomal unconditional driver of male sex chromosomes *A* in a system with no pre-existing sex chromosome drive**

**invasion of *A***: We computed the Jacobian matrix associated with the system of equations (B.9) (with $k_a = k_b = \frac{1}{2}$ and $k_c = k_d = k$), for the equililibrium where the mutant allele $A$ is absent. At this equilibrium, all males are XYaa ($\bar{y}_1 = 1$, $\bar{y}_2 = 0$, $\bar{y}_3 = 0$), and the frequencies of XXaa, XX*aa and X*Yaa females are given by the the eigenvector associated to the leading eigenvalue of transition matrix (A.2) with $k = k^* = \frac{1}{2}$ ($\bar{x}_1 = \frac{1}{2w_{X^*Y}-1}$, $\bar{x}_4 = \bar{x}_7 = \frac{w_{X^*Y}-1}{2w_{X^*Y}-1}$). These values were introduced in the Jacobian matrix and we determined numerically, replacing $w_{X^*Y}$ and $k$ in the Jacobian matrix by different combinations of numerical values (ranging respectively from 1 to 4 (0.02 increment) and 0 to 1 (0.02 increment)), whether the leading eigenvalue associated to the Jacobian is smaller or greater than one. We found that regardless of the value given to $w_{X^*Y}$, the leading eigenvalue of the system is greater than one only if $0.5 < k < 1$. In other words, an autosomal allele which increases the transmission of males' Y chromosomes will increase in frequency when rare in a polygenic sex determination such as the one found in *Mus minutoides*.

**fixation of *A***: Equations (B.9) (with $k_a = k_b = \frac{1}{2}$ and $k_c = k_d = k$) were derived with different numerical combinations of $w_{X^*Y}$ and $k$, ranging respectively from 1 to 4 (0.06 increment) and 0.5 to 1 (0.01 increment). Deterministic simulations were started near the equilibrium with $Y^d$ absent: the initial frequencies of $y_1$, $x_1$, $x_4$ and $x_7$ were set according to the equilibrium frequencies at equilibrium (A.4), and the frequency of $y_2$ was set to $10^{-3}$. Each simulation was run for 500 000 generations or until the change in allele frequency across generations was $< 10^{-20}$. The outcome of these simulations is shown on fig. S2.

Assuming the condition for invasion is respected ($k > 0.5$), a rare autosomal mutant that affects the transmission ratio of male sex chromosome will increase in frequency until it reaches a stable polymorphic equilibrium (see fig. B2, that provides evidence that a stable equilibrium is reached). This frequency is however low across most of the parameter space analyzed. It can reach high frequencies, but only in a limited parameter range, with $k$ very close to 0.5 (the maximum equilibrium frequency reached in our simulation was 0.987, for $k = 0.51$ and $w_{X^*Y} = 1.82$). Our empirical estimation for the value of $k$ in the African pygmy mouse is far from this marginal parameter space: with $k = 0.8$, the maximum equilibrium frequency reached was 0.031 for $w_{X^*Y} = 1.94$. At such low equilibrium frequency, a mutant autosomal driver is unlikely to go to fixation.

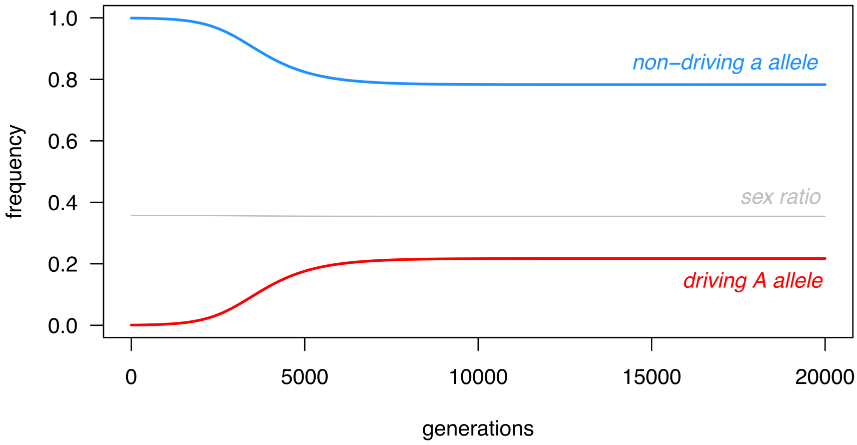

**Figure B2. A Fate of a rare driving *A* in a single simulation.** In this deterministic simulation, $k$ was set to 0.55 and $w_{\text{X*Y}}$ to 2. The simulation was run for 500 000 generations, but only the first 20 000 generations are displayed.

- **step b2': invasion of an autosomal conditional driver of male sex chromosomes *A* in a system with no pre-existing sex chromosome drive**

We proceeded in the same way we did for step b2 to determine the conditions for invasion and fixation of this mutation, with $k_a = k_b = \frac{1}{2}$, $k_c = k$ and $k_d = k*$.

**invasion of *A***: We found that regardless of the numerical value given to $w_{\text{X*Y}}$ and $k*$, the leading eigenvalue of the Jacobian matrix at the equilibrium where $A$ is absent for this system is greater than one only if $0.5 < k < 1$. A rare autosomal allele that skews the transmission of male sex chromosomes in a conditional manner will therefore tend to spread as long as it favours the transmission of the Y in crosses with XX and XX* females, regardless of the transmission of male sex chromosomes in crosses with X*Y females.

**fixation of *A***: The fate of an emergent autosomal mutant that conditionally affects the transmission ratio of male sex chromosomes depends on the three parameters $w_{\text{X*Y}}$, $k$ and $k*$ (fig. S3). If $w_{\text{X*Y}}$ is close to one, a mutant $A$ allele is more likely to go to fixation for values of $k* < 0.5$ and values of $k$ close to 0.5. For larger values of $w_{\text{X*Y}}$ (here 1.5 or 2), $k$ has less impact on the spread of the mutant allele, and the main factor determining if it goes to fixation is whether $k*$ is smaller or greater than roughly 0.5.

Comparing these results to those obtained for step 2b reveals that it is much easier (*i.e.*, possible for greater range of parameters) for a sex chromosome driver carried by an autosome to go to fixation if drive is conditional (fig. S2).

- **Step 3 - invasion of an autosomal driver of male sex chromosomes specific to XY x X*Y crosses in a XX,XX*,X*Y/XY system with pre-existing (unconditional) male sex chromosome drive**

We proceeded in the same way we did for step b2 to determine the conditions for invasion and fixation of this mutation, with $k_a = k_c = k$ and $k_b = k_d = k*$.

**invasion of *A***: for a large range of numerical combinations of $k$ and $k*$, ranging respectively from $\frac{1}{w_{\text{X*Y}}+1}$ (minimum value for the maintenance of the X*, see eq. B.2) to 1 (0.01 increment) and 0 to 1 (0.01 increment), and with $w_{\text{X*Y}}$ taking one of the following values $\{0.5, 1, 2\}$, we found that the system is unstable (leading eigenvalue greater than one) for $k* < k$ *i.e.*, the mutant $A$ allele can invade if it reduces the transmission of male Y chromosome in crosses with X*Y females.

**fixation of *A***: Numerical simulations show that a rare autosomal sex chromosome drive modifier will go to fixation for $k* < k$, condition identical to the condition for invasion (fig. B3).

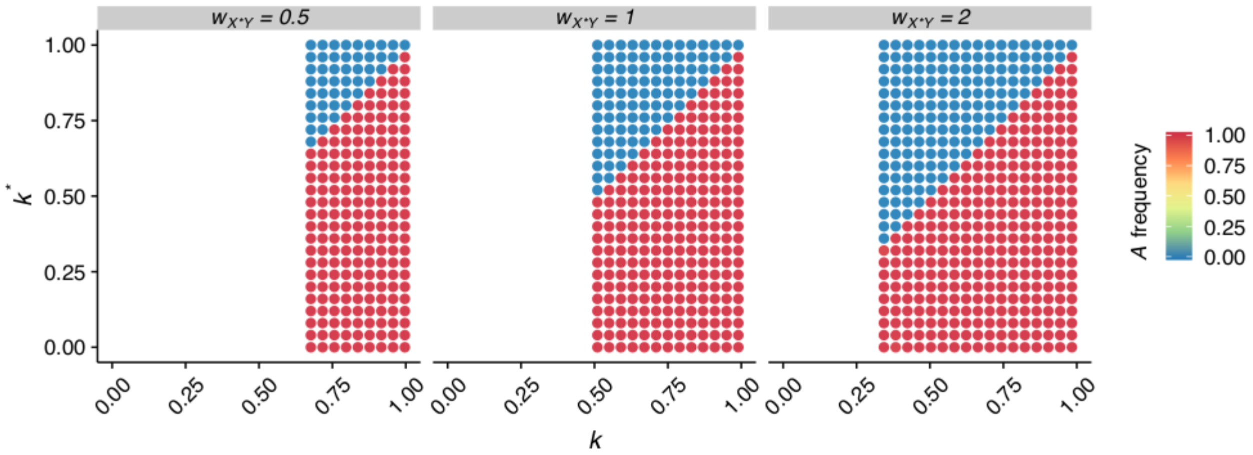

**Figure B3. Fate of a rare autosomal sex chromosome drive modifier in a XX,XX*,X*Y/XY system with pre-existing (unconditional) male sex chromosome drive.** Each dot shows the outcome of a single simulation with fixed values of $k$, $k*$ and $w_{X*Y}$. Simulations were started with a frequency of $A$ set to $10^{-3}$, and ran for 500,000 generations or until the change in frequency of $A$ was $< 10^{-20}$. Color indicates the frequency of the mutant $A$ at the end of the simulation.

### Appendix C - Long term persistence of the polygenic sex determination system: evolution of a suppressor of the X*

In this part, we study the evolution of suppressors of the feminizing activity of the X*, either Y-linked or autosomal. Our aim is to (i) determine the conditions under which a rare suppressor can invade, (ii) explore the consequences of the spread of a suppressor on the sex determination system and (iii) determine how the conditional drive of male sex chromosomes influences these processes.

#### Y-linked suppressor of the X*

##### The model

In a model with X, X*, Y chromosomes, a new allele is introduced: $Y^s$, that suppresses the feminizing activity of the X*, so that $X^*Y^s$ individuals develop as males. A cross between a X*Y female and a $X^*Y^s$ male gives rise to a new female genotype: X*X*. Therefore, there are three types of males: XY, $XY^s$ and $X^*Y^s$ and four types of females: XX, XX*, X*Y and X*X* in numbers $m_j$ ($m_1...m_3$) and $f_i$ ($f_1...f_4$) respectively.

In all females, the transmission ratio of sex chromosomes is Mendelian. In males, the transmission ratio of sex chromosomes is biased: the Y (or $Y^s$) chromosome has a transmission ratio of $k$ in crosses with XX, XX* and X*X* females and $k^*$ in crosses with X*Y females.

We assume that there is no cost for bearing the suppressor allele, but that $X^*Y^s$ males might bear a fertility cost for carrying the X* chromosome ($w_{XY} = w_{XY^s} = 1$ and $w_{X^*Y^s} \leq 1$). As in previous models, X*Y females have a relative fertility of $w_{X^*Y}$. The relative fertility of X*X* females, noted $w_{X^*X^*}$, is also allowed to vary.

The dynamics of the system is given by the following recursions:

$$
\begin{aligned}
{f_1}' &= \frac{1}{\sum_j m_j}\left[(m_1+m_2)(1-k)\left(f_1+\frac{f_2}{2}\right)\right]\\
{f_2}' &= \frac{1}{\sum_j m_j}\left[(m_1+m_2)\left((1-k)\left(\frac{f_2}{2}+f_4 w_{X^*X^*}\right)+\frac{1-k^*}{2}f_3 w_{X^*Y}\right)+m_3 w_{X^*Y^s}(1-k)\left(f_1+\frac{f_2}{2}\right)\right]\\
{f_3}' &= \frac{1}{\sum_j m_j}\left[m_1\left(k\left(\frac{f_2}{2}+f_4 w_{X^*X^*}\right)+\frac{k^*}{2}f_3 w_{X^*Y}\right)+m_3 w_{X^*Y^s}\frac{1-k^*}{2}f_3 w_{X^*Y}\right]\\
{f_4}' &= \frac{1}{\sum_j m_j}\left[m_3 w_{X^*Y^s}\left((1-k)\left(\frac{f_2}{2}+f_4 w_{X^*X^*}\right)+\frac{1-k^*}{2}f_3 w_{X^*Y}\right)\right]\\
{m_1}' &= \frac{1}{\sum_j m_j}\left[m_1\left(k\left(f_1+\frac{f_2}{2}\right)+\frac{1-k^*}{2}f_3 w_{X^*Y}\right)+m_2\frac{1-k^*}{2}f_3 w_{X^*Y}\right]\\
{m_2}' &= \frac{1}{\sum_j m_j}\left[(m_2+m_3 w_{X^*Y^s})\,k\left(f_1+\frac{f_2}{2}\right)\right]\\
{m_3}' &= \frac{1}{\sum_j m_j}\left[(m_2+m_3 w_{X^*Y^s})\left(k\left(\frac{f_2}{2}+f_4 w_{X^*X^*}\right)+\frac{k^*}{2}f_3 w_{X^*Y}\right)\right]
\end{aligned}
\tag{C.1}
$$

where $f_i'$ and $m_j'$ are the number of females and males of genotype $i$ and $j$ at the next generation.

##### Evolutionary stability of the system

Conditions for the evolutionary stability of the XX-XX*-X*Y / XY system (*i.e.*, protection against the spread of a rare $Y^s$) can be derived from equations (C.1), by studying the Jacobian matrix of the system at the equilibrium with $Y^s$ absent. At this equilibrium, all males are XY ($\hat{y}_1 = 1$, $\hat{y}_2 = 0$, $\hat{y}_3 = 0$), and the frequencies of XX, XX* and X*Y females ($\hat{x}_1$, $\hat{x}_2$ and $\hat{x}_3$) are given by the eigenvector of the leading value of the transition matrix of the system (see Appendix A, eq. (A.2)).

The resulting Jacobian matrix is too complex to show here, and did not provide simple analytical eigenvalues. Replacing parameters $k$ and $k^*$ in the matrix by the the empirical values measured in the African pygmy mouse

(0.8 and 0.36), simplified the equations enough to compute these eigenvalues. These values are too complex to show here, but we found that the leading eigenvalue is greater than one under certain circumstances, which provides the conditions for invasion of a Y-linked suppressor of the X*:

$$w_{\mathrm{X^*Y}^s} > \frac{10}{73} \text{ and } w_{\mathrm{X^*Y}} < \frac{80w_{\mathrm{X^*Y}^s} + 275{w_{\mathrm{X^*Y}^s}}^2}{256 - 144w_{\mathrm{X^*Y}^s}} \tag{C.2}$$

The stability of the system depends on the relative fertility of X*Y females and X*Y$^{\mathrm{s}}$ males, and is independent from that of X*X* females (fig. S6). If the cost of bearing an X* in X*Y$^{\mathrm{s}}$ males is too large (if their fertility is less than $\frac{10}{73}$ of that of XY males here), the system is stable, and a rare Y$^{\mathrm{s}}$ cannot invade. Above this threshold, the higher their fertility, the more likely it will invade. The relative fertility of X*Y females has the opposite effect, the higher it is, the more likely the system will resist the spread of the suppressor. If carrying the X* has no impact on the fertility of X*Y$^{\mathrm{s}}$ males ($w_{\mathrm{X^*Y}^s} = 1$), a rare Y$^{\mathrm{s}}$ suppressor will spread unless X*Y females have a fertility more than three times greater than that of XX and XX* females ($w_{\mathrm{X^*Y}} \gtrapprox 3.17$).

In order to explore the impact of $k$ and $k^*$ on the stability of the system, we performed a numerical analysis of the model. For a large range of numerical combinations of $w_{\mathrm{X^*Y}}$, $w_{\mathrm{X^*Y}^s}$, $k$ and $k^*$, we determined whether the leading eigenvalue associated to the Jacobian is smaller or greater than one ($w_{\mathrm{X^*X^*}}$ was set to one, as it has no impact on stability; eq. (C.2)). Results are shown on fig. S4: the value of $k$ has little impact on the spread of a suppressor, while high values of $k^*$ tend to promote the spread of a Y$^{\mathrm{s}}$. Overall, evolutionary stability is much more likely if $k^* < 0.5$ *i.e.*, if males see their X chromosome transmitted more often in crosses with X*Y females, as observed in our empirical case study.

#### Consequences of the spread of a Y-linked suppressor of X* activity

Following the spread of a Y-linked suppressor of the X* (Y$^{\mathrm{s}}$), different equilibria could theoretically be reached depending on which alleles remain in the system. All possibilities and paths to the different equilibria are displayed on fig. C1. In particular, simple heterogamety could be restored, with either (i) X*X* females and X*Y$^{\mathrm{s}}$ males, assuming the X and the non-suppressor Y are lost (fig. C1, red equilibrium), or (ii) XX females and XY$^{\mathrm{s}}$ (and/or XY) males (yellow and orange equilibria), if the X* is lost. Alternatively, a stable "alternative" polygenic equilibrium could be reached, with multiple sex chromosome combinations segregating within at least one sex. This is true if (i) all four alleles are maintained (X, X*, Y and Y$^{\mathrm{s}}$, dark blue equilibrium), (ii) the X is lost alone (light blue), or (iii) the non-suppressor Y is lost alone (purple).

Deterministic numerical simulations were performed to define which equilibria could be reached, and under which conditions a return to standard heterogamety is possible. The recursion equations (C.1) were derived with different combinations of $w_{\mathrm{X^*Y}}$ and $w_{\mathrm{X^*Y}^s}$ ranging from respectively from 0 to 2 (0.08 increment) and 0 to 1 (0.04 increment). The fertility of X*X* females was set to either (i) $w_{\mathrm{X^*X^*}} = 1$, or (ii) $w_{\mathrm{X^*X^*}} = w_{\mathrm{X^*Y}}$ (*i.e.*, same fertility as XX and XX* females or as X*Y females). These two cases make the most sense biologically considering the difference in reproductive success between XX and XX* females versus X*Y females. Simulations were started near the equilibrium with $Y^s$ absent: the initial frequencies $y_1$, $x_1$, $x_2$ and $x_3$ were set according to the equilibrium frequencies obtained in Appendix A (eq. (A.4)), and $y_2$ was set to $10^{-3}$. Each simulation was run for 50'000 generations or until the change in allele frequency was $< 10^{-20}$. At the end of each simulation, the presence or absence of all four alleles (X, X*, Y and Y$^{\mathrm{s}}$) was assessed, assuming an allele was absent if its frequency was below 0.001. Results of these simulations are displayed on the upper row of fig. C2.

If the conditions for invasion of the suppressor are met (eq. C.2), different equilibria can indeed be reached, depending on the values of $w_{\mathrm{X^*Y}}$, $w_{\mathrm{X^*Y}^s}$ and $w_{\mathrm{X^*X^*}}$. Both the Y and Y$^{\mathrm{s}}$ were maintained across most of the parameter space explored (dark blue, light blue and yellow areas). Depending on the fertility of X*Y females (mostly), one of the two X chromosomes can be lost: the X chromosome was lost for higher fertility of X*Y females (light blue), while the X* was lost if their fertility is too low (yellow and orange). Higher fertility of X*Y$^{\mathrm{s}}$ males (close to one) also tended to favour the loss of the X*. Finally, the non-suppressor Y allele was only lost if X*Y$^{\mathrm{s}}$ males bear no or very little cost for carrying the X*, resulting in a return to standard male heterogamety with XX and XY$^{\mathrm{s}}$ males (orange area). The equilibrium where the ancestral Y is lost, and the X, X* and Y$^{\mathrm{s}}$ are maintained at equilibrium (purple area), was only reached with $w_{\mathrm{X^*X^*}} = 1$. This equilibrium is only neutrally stable, as in these conditions, all female genotypes (XX, XX* and X*X*) and male genotypes (XY$^{\mathrm{s}}$ and X*Y$^{\mathrm{s}}$) present at equilibrium have the same fertility. Stochastic processes, or a slight difference in fitness among these genotypes (*e.g.*, a small cost for bearing the suppressor in X*Y$^{\mathrm{s}}$), will eventually lead to the loss of either the X or the X*, causing a return to a standard male heterogametic system (red or orange area). The same goes with the yellow equilibrium, in which all females are XX and males either XY or XY$^{\mathrm{s}}$: either the Y or Y$^{\mathrm{s}}$ is eventually bound to disappear because of genetic drift or fitness differences.

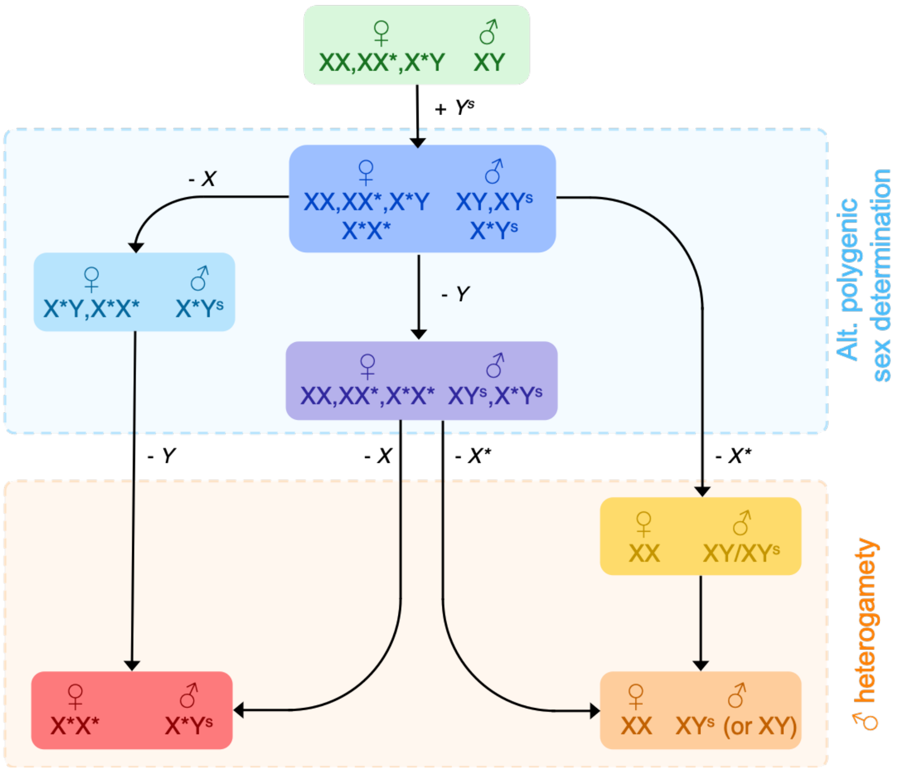

**Figure C1. Paths leading to heterogamety following the spread of a Y-linked suppressor of X* activity.** Each solid square represents a putative stable equilibrium, with the genotype of females (left) and males (right) present at that equilibrium. Along each branch leading from one equilibrium to the next is displayed which allele is lost (or gained in the first step). The three equilibria with cold colors (shaded blue area) are those where multiple genotypes co-exist in at least one sex (polygenic sex determination), and the three with warm colors (shaded orange area) are those where sex determination corresponds to a standard heterogametic system

To verify this we re-ran simulation in which we slightly modified the fertility of $XY^s$ males (+/-0.01) (fig. C2, middle and bottom rows). In both cases, the two "neutrally stable" equilibria were not reached. With $w_{X^*X^*} = 1$, in the parameter space that previously reached the purple equilibrium, the X chromosome was lost if $XY^s$ males had a slight fitness disadvantage (middle left plot) and the X* was lost if they had a slight fitness advantage (lower left plot). With both $w_{X^*X^*} = 1$ and $w_{X^*X^*} = w_{X^*Y}$, in the parameter space that previously reached the yellow equilibrium, if $XY^s$ had a slight fitness disadvantage (middle row), the X* was not eliminated, and if they had a slight fitness advantage (bottom row), both the X* and the non-suppressor Y were lost.

#### Influence of the conditional nature of male sex chromosome drive

fig. S4 shows that male sex chromosome drive has an impact on the spread of a rare Y-linked suppressor of X* activity. In order to further evaluate how the conditional nature of the drive observed in *Mus minutoides* influences the conditions under which the spread of such a suppressor could lead to a return to standard male heterogamety, we ran additional simulations based on recursion equations (C.1), setting the values of $k$ an $k^*$ to $k = k^* = 0.8$ (to emulate a non-conditional drive) or $k = k^* = 0.5$ (absence of drive). Results are displayed on fig. C3, alongside the results obtained for $k = 0.8$ and $k^* = 0.36$, already shown on fig. C2.

With both unconditional drive and no drive, a Y-linked suppressor is more likely to spread compared to when drive is conditional (fig. C3, reduction of the green area). The setting the most vulnerable to invasion by a mutant suppressor is the unconditional setting (fig. C3b,e), in line with our previous results demonstrating that high values of $k^*$ make the system more vulnerable (fig. S4). Furthermore, in both cases (unconditional drive or no drive), a suppressor that spreads in the population is also much more likely to cause a return to a standard male heterogametic system than in the presence of conditional drive, as shown on fig. C3 by the increased size of the red and orange areas, as well as the purple and yellow ones (that as explained earlier, are only neutrally stable, and will eventually lead the system to revert to male heterogamety). The impact of $XY^s$ male fertility $w_{XY^s}$ is similar whether drive is conditional, unconditional or absent: the closest it is to 1, the more likely the spread of a suppressor will lead to the loss of the non-suppressor Y and cause a return to standard male heterogamety. Higher fertility of X*Y females raises the chance that a stable polymorphic equilibrium is reached (blue area), but only if the fertility of X*X* females is equivalent to that of XX and XX* females (upper plots on fig. C3). If X*X* females have a similar fertility to X*Y females (lower plots), the probability of a transition to a X*X*/X*Y$^s$ (red areas) increases with their fertility, but only if drive is unconditional or absent.

To summarise, in addition to increasing the stability of the system against the spread of a Y-linked suppressor of the X*, the conditional nature of male sex chromosome drive found in the African pygmy mouse also reduces the likelihood that the polygenic sex determination system reverts back to a standard male heterogamety system under the influence of such a suppressor.

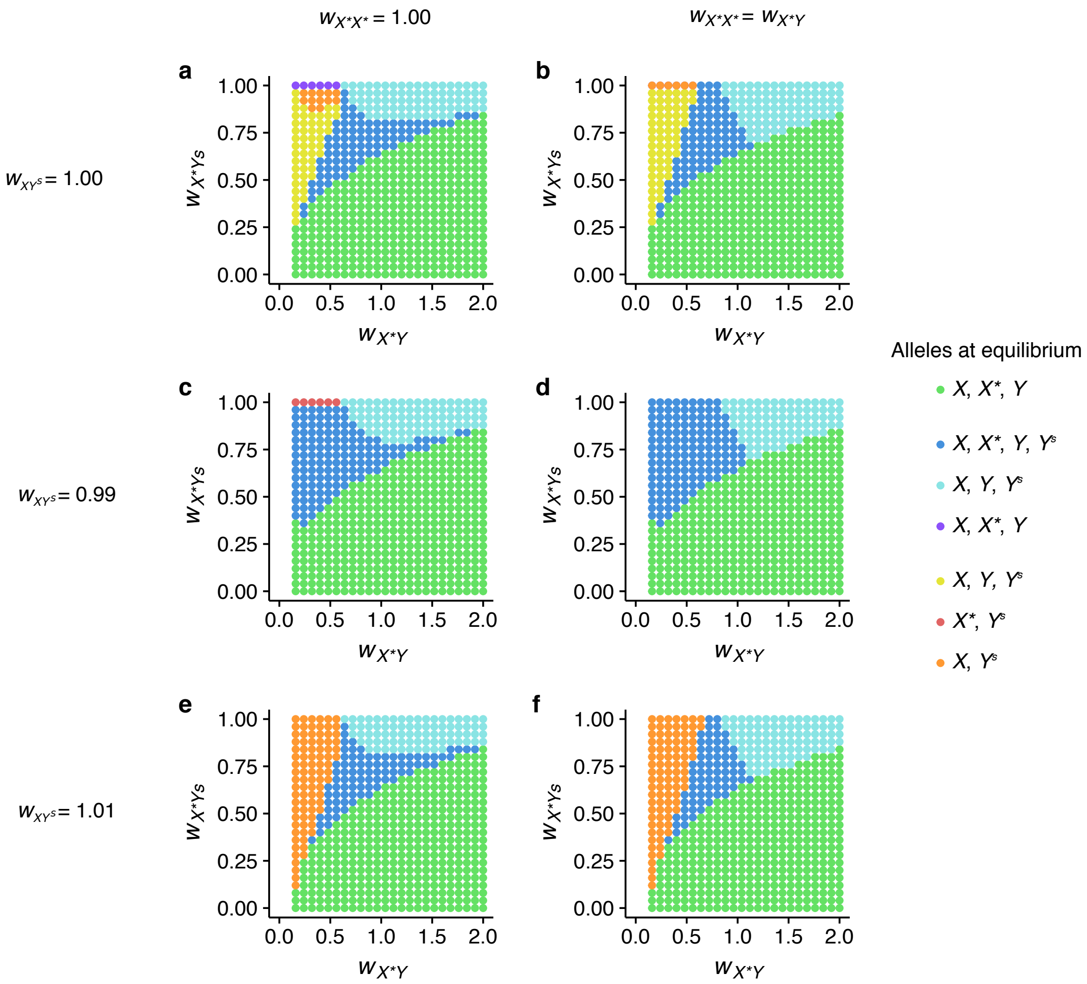

**Figure C2. Fate of a rare Y-linked suppressor of X* feminizing activity (Y$^s$) in a XX,XX*,X*Y/XY system.** $k$ and $k^*$ were set to 0.8 and 0.36 to mimic the conditional drive of male sex chromosomes found in the African pygmy mouse. Each individual plot corresponds to a combination of fixed values for $w_{X^*X^*}$ (columns) and $w_{XY^s}$ (rows). Within each plot, each dot shows the outcome of a single deterministic simulation with fixed values of $w_{X^*Y}$ and $w_{X^*Y^s}$. Different colors indicate the equilibrium reached at the end of a simulation (see fig. C1). White areas corresponds to the parameter space which does not allow maintenance of the X* (eq. (1) in main text). The comparison of simulations with $w_{XY^s}$ =1, 0.99 or 1.01 illustrates that some equilibria (purple and yellow areas) are only neutrally stable in the parameter space initially explored, as a very slight change in the fertility of XY$^s$ males render them unstable.

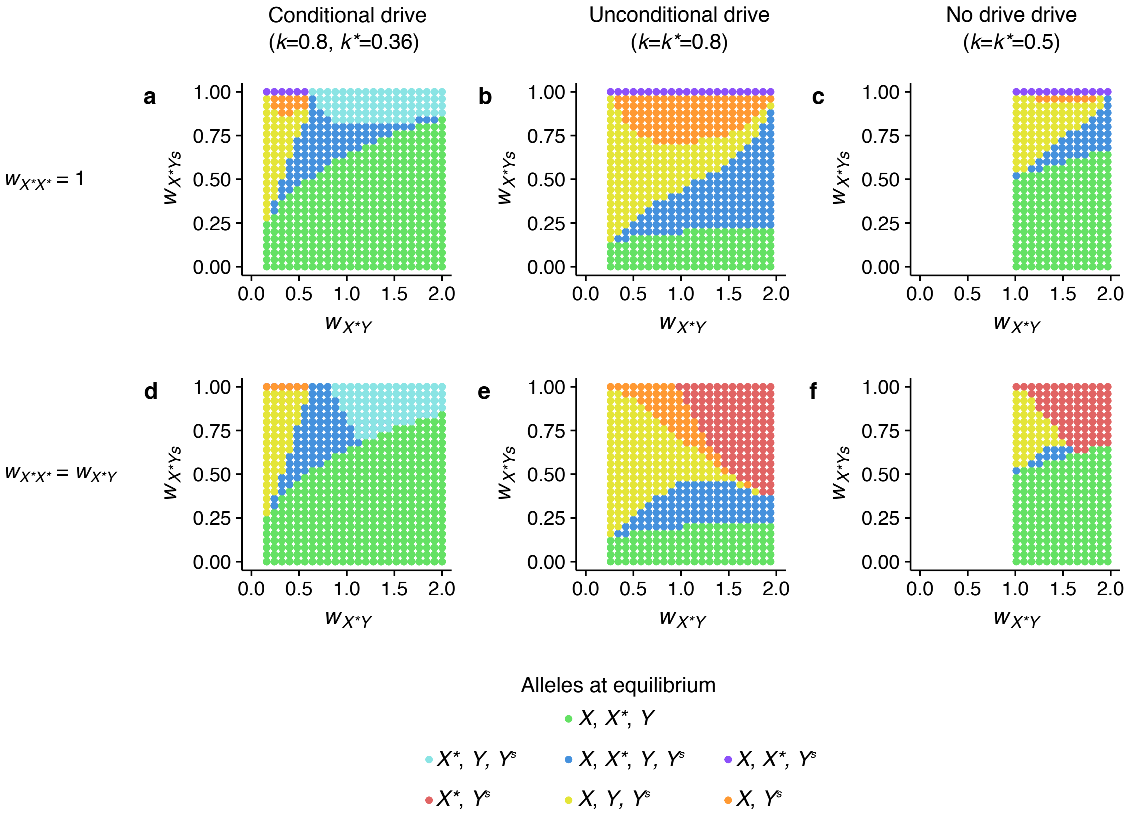

**Figure C3. Fate of a rare Y-linked suppressor of X* feminizing activity ($Y^s$) in a XX,XX*,X*Y/XY system**. With either conditional drive ($k = 0.8$, $k^* = 0.36$), unconditional drive ($k = 0.8$, $k^* = 0.8$), or no sex chromosome drive ($k = 0.5$, $k^* = 0.5$) (columns), and different values for $w_{X^*X^*}$ (rows). Each dot shows the outcome of a single deterministic simulation with fixed values of $w_{X^*Y}$ and $w_{X^*Y^s}$. Different colors indicate the equilibrium reached at the end of a simulation (see fig. C1). White areas corresponds to the parameter space which does not allow maintenance of the X* (eq. (1) in main text). The two plots displayed in the left column (conditional drive), are identical to those displayed in the upper row of fig. C2.

### Autosomal suppressor of the X*

#### The model

At the sex determining locus, the alleles considered are: Y, X and X*. An autosomal locus is added to the model, with two alleles: the ancestral $a$ allele, and a dominant mutant allele $A$, which suppresses the feminizing action of the X* chromosome, and turns X*YAa and X*YAA individuals into males. A cross between a X*Y male and X*Y female gives rise to X*X* females. Therefore, there are five types of males: XYaa, XYAa, XYAA, X*YAa and X*YAA and ten types of females: XXaa, XXAa, XXAA, XX*aa, XX*Aa, XX*AA, X*Yaa, X*X*aa, X*X*Aa and X*X*AA, in numbers $m_j$ ($m_1 ... m_5$) and $f_i$ ($f_1 ... f_{10}$) respectively.

In all females, the transmission ratio of sex chromosomes is Mendelian. In males, the transmission ratio of sex chromosomes is biased: the Y chromosome has a transmission ratio of $k$ in crosses with XX, XX* and X*X* females and $k^*$ in crosses with X*Y females (regardless of genotype at the autosomal locus).

We assume that the autosomal locus has no impact on fitness, but that X*Y males might bear a fertility cost for carrying the X* chromosome, their relative fertility is noted: $w_{X^*Y♂}$ ($\leq 1$). As in previous models, XX and XX* females are assumed to have a similar fertility, X*Y and X*X* females have a relative fertility of $w_{X^*Y♀}$ and $w_{X^*X^*}$ respectively.

The dynamics of the system is given by the following recursions:

$$m_1' = \frac{1}{\sum_j m_j}\left[\left(m_1 + \frac{m_2}{2}\right)\left(k\left(f_1 + \frac{f_2}{2} + \frac{f_4}{2} + \frac{f_5}{4}\right) + \frac{1-k^*}{2} f_7 w_{X^*Y♀}\right) + m_4 w_{X^*Y♂}\frac{k}{2}\left(f_1 + \frac{f_2}{2} + \frac{f_4}{2} + \frac{f_5}{4}\right)\right]$$

$$m_2' = \frac{1}{\sum_j m_j}\left[m_1 k\left(\frac{f_2}{2} + f_3 + \frac{f_5}{4} + \frac{f_6}{2}\right) + m_2\left(k\left(\frac{f_1+f_2+f_3}{2} + \frac{f_4+f_5+f_6}{4}\right) + \frac{1-k^*}{4} f_7 w_{X^*Y♀}\right) + m_3\left(k\left(f_1 + \frac{f_2}{2} + \frac{f_4}{2} + \frac{f_5}{4}\right) + \frac{1-k^*}{2} f_7 w_{X^*Y♀}\right) + m_4 w_{X^*Y♂} k\left(\frac{f_1+f_2+f_3}{2} + \frac{f_4+f_5+f_6}{4}\right) + m_5 w_{X^*Y♂} k\left(f_1 + \frac{f_2}{2} + \frac{f_4}{2} + \frac{f_5}{4}\right)\right]$$

$$m_3' = \frac{1}{\sum_j m_j}\left[\left(\frac{m_2}{2} + m_3 + \left(\frac{m_4}{2} + m_5\right) w_{X^*Y♂}\right) k\left(\frac{f_2}{2} + f_3 + \frac{f_5}{4} + \frac{f_6}{2}\right)\right]$$

$$m_4' = \frac{1}{\sum_j m_j}\left[m_1 k\left(\frac{f_5}{4} + \frac{f_6}{2} + \left(\frac{f_9}{2} + f_{10}\right) w_{X^*X^*}\right) + (m_2 + m_4 w_{X^*Y♂})\, k\left(\frac{f_4+f_5+f_6}{4} + \frac{f_8+f_9+f_{10}}{2} w_{X^*X^*}\right) + (m_3 + m_5 w_{X^*Y♂}) k\left(\frac{f_4}{2} + \frac{f_5}{4} + \left(f_8 + \frac{f_9}{2}\right) w_{X^*X^*}\right) + \left(\left(\frac{m_2}{2} + m_3\right)\frac{k^*}{2} + \left(\frac{m_4}{4} + \frac{m_5}{2}\right) w_{X^*Y♂}\right) f_7 w_{X^*Y♀}\right]$$

$$m_5' = \frac{1}{\sum_j m_j}\left[\left(\frac{m_2}{2} + m_3 + \left(\frac{m_4}{2} + m_5\right) w_{X^*Y♂}\right) k\left(\frac{f_5}{4} + \frac{f_6}{2} + \left(\frac{f_9}{2} + f_{10}\right) w_{X^*X^*}\right)\right]$$

$$f_1' = \frac{1}{\sum_j m_j}\left[\left(m_1 + \frac{m_2}{2}\right)(1-k)\left(f_1 + \frac{f_2}{2} + \frac{f_4}{2} + \frac{f_5}{4}\right)\right]$$

$$f_2' = \frac{1}{\sum_j m_j}\left[m_1(1-k)\left(\frac{f_2}{2} + f_3 + \frac{f_5}{4} + \frac{f_6}{2}\right) + m_2(1-k)\left(\frac{f_1+f_2+f_3}{2} + \frac{f_4+f_5+f_6}{4}\right) + m_3(1-k)\left(f_1 + \frac{f_2}{2} + \frac{f_4}{2} + \frac{f_5}{4}\right)\right]$$

$$f_3' = \frac{1}{\sum_j m_j}\left[\left(\frac{m_2}{2} + m_3\right)(1-k)\left(\frac{f_2}{2} + f_3 + \frac{f_5}{4} + \frac{f_6}{2}\right)\right]$$

$$
\begin{aligned}
f_4{}' &= \frac{1}{\sum_j m_j}\Bigg[\left(m_1+\frac{m_2}{2}\right)\left((1-k)\left(\frac{f_4}{2}+\frac{f_5}{4}+\left(f_8+\frac{f_9}{2}\right)w_{\mathrm{X^*X^*}}\right)+\frac{1-k^*}{2}f_7 w_{\mathrm{X^*Y}\female}\right)+\\
&\qquad m_4 w_{\mathrm{X^*Y}\male}\frac{1-k}{2}\left(f_1+\frac{f_2}{2}+\frac{f_4}{2}+\frac{f_5}{4}\right)\Bigg]\\
f_5{}' &= \frac{1}{\sum_j m_j}\Bigg[m_1(1-k)\left(\frac{f_5}{4}+\frac{f_6}{2}+\left(\frac{f_9}{2}+f_{10}\right)w_{\mathrm{X^*X^*}}\right)+\\
&\qquad m_2\left((1-k)\left(\frac{f_4+f_5+f_6}{4}+\frac{f_8+f_9+f_{10}}{2}w_{\mathrm{X^*X^*}}\right)+\frac{1-k^*}{4}f_7 w_{\mathrm{X^*Y}\female}\right)+\\
&\qquad m_3\left((1-k)\left(\frac{f_4}{2}+\frac{f_5}{4}+\left(f_8+\frac{f_9}{2}\right)w_{\mathrm{X^*X^*}}\right)+\frac{1-k^*}{2}f_7 w_{\mathrm{X^*Y}\female}\right)+\\
&\qquad m_4 w_{\mathrm{X^*Y}\male}(1-k)\left(\frac{f_1+f_2+f_3}{2}+\frac{f_4+f_5+f_6}{4}\right)+m_5 w_{\mathrm{X^*Y}\male}(1-k)\left(f_1+\frac{f_2}{2}+\frac{f_4}{2}+\frac{f_5}{4}\right)\Bigg]\\
f_6{}' &= \frac{1}{\sum_j m_j}\Bigg[\left(\frac{m_2}{2}+m_3\right)(1-k)\left(\frac{f_5}{4}+\frac{f_6}{2}+\left(\frac{f_9}{2}+f_{10}\right)w_{\mathrm{X^*X^*}}\right)+\\
&\qquad \left(\frac{m_4}{2}+m_5\right)w_{\mathrm{X^*Y}\male}(1-k)\left(\frac{f_2}{2}+f_3+\frac{f_5}{4}+\frac{f_6}{2}\right)\Bigg]\\
f_7{}' &= \frac{1}{\sum_j m_j}\Bigg[\left(m_1+\frac{m_2}{2}+\frac{m_4}{2}w_{\mathrm{X^*Y}\male}\right)k\left(\frac{f_4}{2}+\frac{f_5}{4}+\left(f_8+\frac{f_9}{2}\right)w_{\mathrm{X^*X^*}}\right)+\\
&\qquad \left(m_1+\frac{m_2}{2}\right)\frac{k^*}{2}f_7 w_{\mathrm{X^*Y}\female}+m_4 w_{\mathrm{X^*Y}\male}\frac{f_7}{4}w_{\mathrm{X^*Y}\female}\Bigg]\\
f_8{}' &= \frac{1}{\sum_j m_j}\Bigg[m_4 w_{\mathrm{X^*Y}\male}\left(\frac{1-k}{2}\left(\frac{f_4}{2}+\frac{f_5}{4}+\left(f_8+\frac{f_9}{2}\right)w_{\mathrm{X^*X^*}}\right)+\frac{1-k^*}{2}f_7 w_{\mathrm{X^*Y}\female}\right)\Bigg]\\
f_9{}' &= \frac{1}{\sum_j m_j}\Bigg[m_4 w_{\mathrm{X^*Y}\male}\left((1-k)\left(\frac{f_4+f_5+f_6}{4}+\frac{f_8+f_9+f_{10}}{2}w_{\mathrm{X^*X^*}}\right)+\frac{1-k^*}{4}f_7 w_{\mathrm{X^*Y}\female}\right)+\\
&\qquad m_5 w_{\mathrm{X^*Y}\male}\left((1-k)\left(\frac{f_4}{2}+\frac{f_5}{4}+\left(f_8+\frac{f_9}{2}\right)w_{\mathrm{X^*X^*}}\right)+\frac{1-k^*}{2}f_7 w_{\mathrm{X^*Y}\female}\right)\Bigg]\\
f_{10}{}' &= \frac{1}{\sum_j m_j}\Bigg[\left(\frac{m_4}{2}+m_5\right)w_{\mathrm{X^*Y}\male}(1-k)\left(\frac{f_5}{4}+\frac{f_6}{2}+\left(\frac{f_9}{2}+f_{10}\right)w_{\mathrm{X^*X^*}}\right)\Bigg]
\end{aligned}
\tag{C.3}
$$

where $f_i'$ and $m_j'$ are the number of females and males of genotype $i$ and $j$ at the next generation.

#### Evolutionary stability of the system

Analytical conditions for the evolutionary stability of the XX-XX*-X*Y / XY system (*i.e.*, protection against the spread of a rare autosomal suppressor) can be derived from equations (C.3), by studying the Jacobian matrix of the system at an equilibrium with $A$ absent. At this equilibrium, all males are XYaa ($\hat{y}_1 = 1$) and the frequencies of XXaa, XX*aa and X*Yaa females ($\hat{x}_1$, $\hat{x}_4$ and $\hat{x}_7$) are given by the eigenvector of the leading value of the transition matrix of the system (see Appendix A, eq. (A.2)).

The resulting Jacobian matrix is too complex to show here, and did not provide simple analytical eigenvalues. We therefore performed a numerical analysis of the system. We first replaced parameters $k$ and $k^*$ in the matrix by the the empirical values measured in the African pygmy mouse (0.8 and 0.36). For a large range of numerical combinations of $w_{\mathrm{X^*Y}\female}$, $w_{\mathrm{X^*Y}\male}$, and $w_{\mathrm{X^*X^*}}$, we determined whether the leading eigenvalue associated to the Jacobian is smaller or greater than one, which gives the conditions under which a rare autosomal suppressor of the feminizing activity of the X* can invade. Like for the Y-linked suppressor, the spread of an autosomal suppressor becomes more likely with increasing values of X*Y male fertility and decreasing values of X*Y female fertility, but additionally, its spread is also impacted by the fertility of X*X* females: increasing values of $w_{\mathrm{X^*X^*}}$ also tend to favour invasion (fig. S6)). The impact of $k$ and $k^*$ on the stability of the system was explored by performing further numerical analyses of the model for a large range of numerical combinations of $w_{\mathrm{X^*Y}\female}$, $w_{\mathrm{X^*Y}\male}$, $k$ and $k^*$. The fertility of X*X* females was set to either (i) $w_{\mathrm{X^*X^*}} = 1$, or (ii) $w_{\mathrm{X^*X^*}} = w_{\mathrm{X^*Y}\female}$ (*i.e.*, same fertility as XX and XX* females or as X*Y females). Results are shown on fig. S5. Alike for a Y suppressor (fig. S4), $k^*$ has the most impact on the stability of the system, with decreasing values of $k^*$ making the spread of an autosomal suppressor less likely.

#### Consequences of the spread of an autosomal suppressor of X* activity

fig. C4 displays all possible equilibra that can be reached following the spread of an autosomal suppressor $A$.

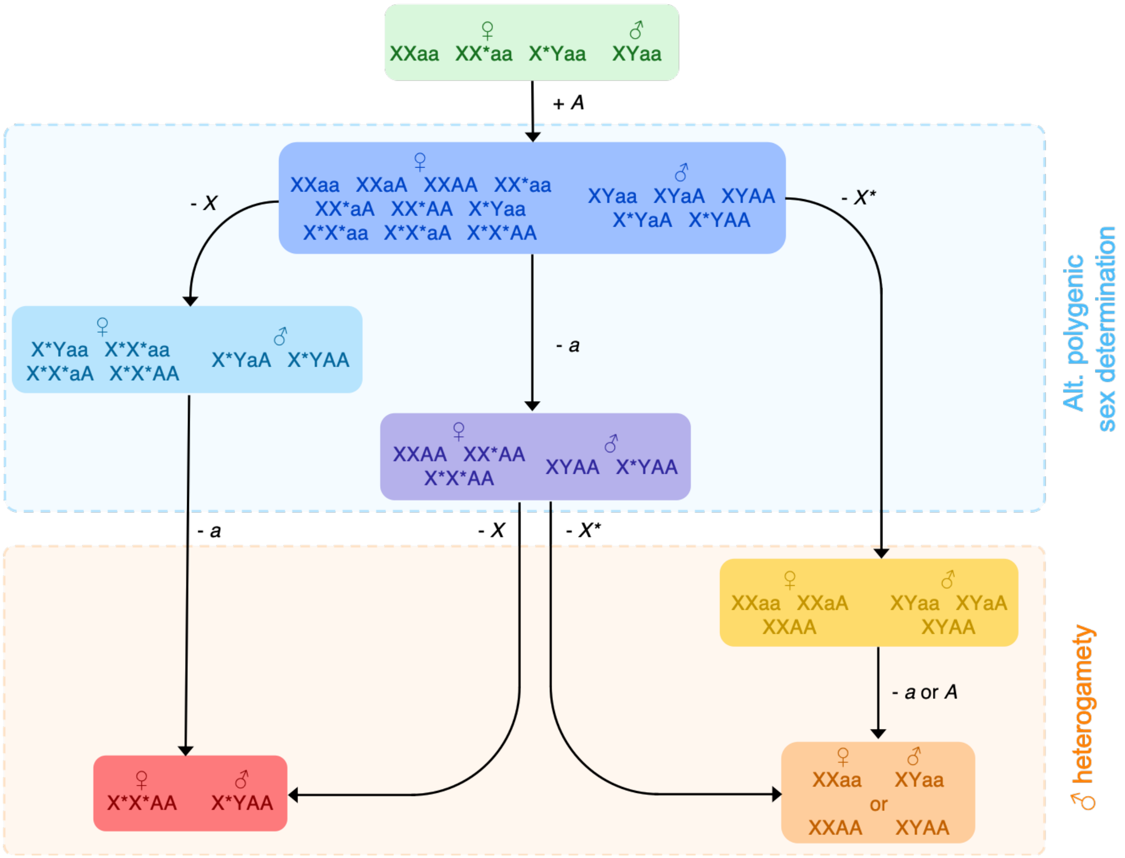

**Figure C4. Paths leading to heterogamety following the spread of an autosomal suppressor of X* activity.** Each solid square represents a putative stable equilibrium, with the genotype of females (left) and males (right) present at that equilibrium. Along each branch leading from one equilibrium to the next is displayed which allele is lost (or gained in the first step). The three equilibria with cold colors (shaded blue area) are those where multiple genotypes co-exist in at least one sex (polygenic sex determination), and the three with warm colors (shaded orange area) are those where sex determination corresponds to a standard heterogametic system.

A stable "alternative" polygenic equilibrium could be reached, with multiple sex chromosome combinations segregating within at least one sex. This is true if (i) all three alleles at the sex determining locus, and the two alleles at the autosomal locus are maintained (dark blue equilibrium), (ii) the X is lost alone (light blue), or (iii) the non-suppressor $a$ allele is lost alone (purple). Simple heterogamety could be also restored, with either (i) X*X* females and X*Y males, following the loss of the X and $a$ (red equilibrium), or (ii) XX females and XY males, following the loss of the X* (yellow) or the X* and either the $a$ or $A$ allele (orange).

We ran deterministic numerical simulations to define which equilibria could be reached following the spread of an autosomal suppressor of the feminizing activity of the X*, and under which conditions a return to standard heterogamety occurs. The recursion equations (C.3) were derived with different numerical combinations of $w_{X^*Y♀}$ and $w_{X^*Y♂}$ ranging from respectively from 0 to 2 (0.08 increment) and 0 to 1 (0.04 increment). The fertility of X*X* females was set to either $w_{X^*X^*} = 1$ or $w_{X^*X^*} = w_{X^*Y♀}$. Finally, to evaluate the impact of the conditional drive of male sex chromosomes, $k$ and $k^*$ were set either to 0.8 and 0.36 respectively (to mimic the conditional drive found in the African pygmy mouse), to $k = k^* = 0.8$ (to emulate a non-conditional drive) or $k = k^* = 0.5$ (absence of drive). Simulations were started near the equilibrium with $A$ absent: the initial frequencies of XYaa males ($y_1$), XXaa, XX*aa and X*Yaa females ($x_1$, $x_4$ and $x_7$) were set according to the equilibrium frequencies obtained in Appendix A (eq. (A.4)), and frequency of X*YaA males ($y_4$) was set to $10^{-3}$. Each simulation was run for 500 000 generations or until the change in allele frequency across generations was $< 10^{-20}$. At the end of each simulation, the presence or absence of all alleles (X, X*, Y, $a$ and $A$) was assessed, assuming an allele was absent if its frequency was below 0.001.

As with the Y-linked suppressor, different equilibria can be reached following the spread of an autosomal suppressor (fig. C5). In our simulations, sex determination remained polygenic across most of the parameter space explored (dark blue and light blue areas), the X being lost for high values of X*Y female and male fertility ($w_{X^*Y♀}$, $w_{X^*Y♂}$, light blue). A return to simple male heterogamety occurred with the X* being lost, but only for low values of X*Y females fertility (yellow area, which is only neutrally stable in the absence of a cost for carrying the suppressor: as all individuals have the same fertility, polymorphism at the autosomal locus will ultimately be lost). In contrast with simulations of a Y-linked suppressor, a X*X*/X*Y equilibrium was never reached. However, note that if $w_{X^*X^*} = w_{X^*Y♀}$, the equilibrium reached following the loss of the X (light blue) is also neutrally stable in the absence of a cost for carrying the suppressor allele $A$. As $A$ cannot be lost, because no males are homozygous for the $a$ allele at this equilibrium, genetic drift will eventually lead to the loss of $a$, and fixation of a X*X*AA/X*YAA heterogametic system.

Concerning the influence of male sex chromosome drive, just like for a Y-linked suppressor, with no drive and especially unconditional drive of the Y, an autosomal suppressor is more likely to spread when rare compared to when drive is conditional (fig. C5, reduction of the green area). This is in line with our results demonstrating that high values of $k^*$ make the system more vulnerable to suppressors (fig. S5, fig. S6). The difference between the three settings is less striking than with a Y-linked suppressor (see fig. C3). Regarding the potential for a return to standard male heterogamety, conditional drive with $k = 0.8$ and $k^* = 0.36$, reduced the parameter space in which it occurred in our simulations (yellow area), but only very slightly.

Overall, despite being possible, the invasion of an autosomal suppressor of the feminizing activity is unlikely to cause a return to male heterogamety, and less so than a Y suppressor.

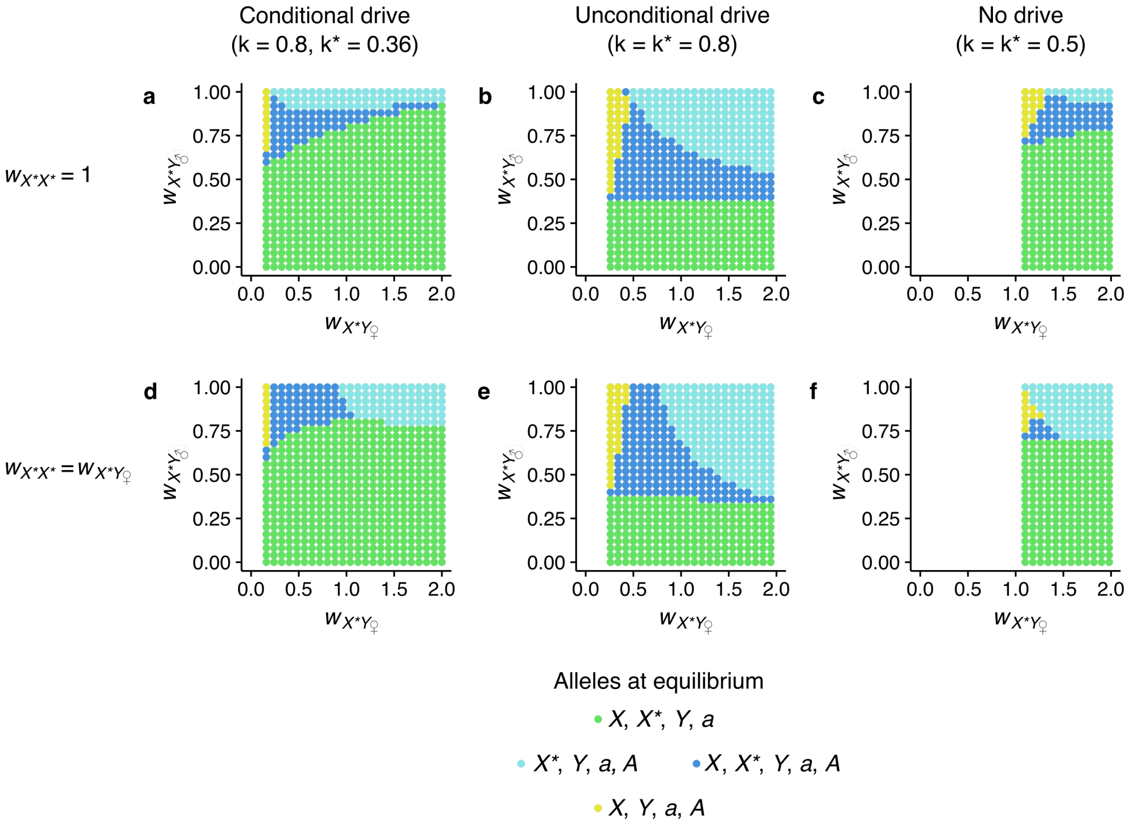

**Figure C5. Fate of a rare autosomal suppressor of X* feminizing activity (Y$^s$) in a XX,XX*,X*Y/XY system**. With either conditional drive ($k = 0.8$, $k^* = 0.36$), unconditional drive ($k = 0.8$, $k^* = 0.8$), or no sex chromosome drive ($k = 0.5$, $k^* = 0.5$) (columns), and different values for $w_{X^*X^*}$ (rows). Each dot shows the outcome of a single deterministic simulation with fixed values of $w_{X^*Y♀}$ and $w_{X^*Y♂}$. Different colors indicate the equilibrium reached at the end of a simulation (see fig. C4). White areas corresponds to the parameter space which does not allow maintenance of the X* (eq. (1) in main text).
